# Reproductive Adaptation of *Astyanax mexicanus* Under Nutrient Limitation

**DOI:** 10.1101/2025.02.13.638191

**Authors:** Fanning Xia, Ana Santacruz, Di Wu, Sylvain Bertho, Elizabeth Fritz, Pedro Morales-Sosa, Sean McKinney, Stephanie H. Nowotarski, Nicolas Rohner

## Abstract

Reproduction is a fundamental biological process for the survival and continuity of species. Examining changes in reproductive strategies offers valuable insights into how animals have adapted their life histories to different environments. Since reproduction is one of the most energy-intensive processes in female animals, nutrient scarcity is expected to interfere with the ability to invest in gametes. Lately, a new model to study adaptation to nutrient limitation has emerged; the Mexican tetra *Astyanax mexicanus*. This fish species exists as two different morphs, a surface river morph and a cave-dwelling morph. The cave-dwelling morph has adapted to the dark, biodiversity, and nutrient-limited cave environment and consequently evolved an impressive starvation resistance. However, how reproductive strategies have adapted to nutrient limitations in this species remains poorly understood. Here, we compared breeding activities and maternal contributions between laboratory-raised surface fish and cavefish. We found that cavefish produce different clutch sizes of eggs with larger yolk compared to surface fish, indicating a greater maternal nutrient deposition in cavefish embryos. To systematically characterize yolk compositions, we used untargeted proteomics and lipidomics approaches to analyze protein and lipid profiles in 2-cell stage embryos and found an increased proportion of sphingolipids in cavefish compared to surface fish. Additionally, we generated transcriptomic profiles of surface fish and cavefish ovaries using a combination of single cell and bulk RNA sequencing to examine differences in maternal contribution. We found that genes essential for hormone regulation were upregulated in cavefish follicular somatic cells compared to surface fish. To evaluate whether these differences contribute to their reproductive abilities under natural-occurring stress, we induced breeding in starved female fish. Remarkably, cavefish maintained their ability to breed under starvation, whereas surface fish largely lost this ability. We identified *insulin-like growth factor 1a receptor* (*igf1ra*) as a potential candidate gene mediating the downregulation of ovarian development genes, potentially contributing to the starvation-resistant fertility of cavefish. Taken together, we investigated the female reproductive strategies in *Astyanax mexicanus*, which will provide fundamental insights into the adaptations of animals to environments with extreme nutrient deficit.

## Introduction

Species survival relies on reproductive success. Nutritional changes in the environment can affect species continuity by interfering with female reproductive capacity^3,4^, as reproduction consumes significant amounts of energy in female animals^5–7^. Human population studies from historical famine events such as Dutch hunger winter, suggested that food deprivation impacts not only the exposed individuals but also has transgenerational effects on their offsprings^8^. Similarly, studies in lab model animals have shown that maternal resource limitation affects both reproductive traits and development in progeny^9–12^.

Despite these challenges, environmental changes do not always lead to extinction. Many species have evolved strategies to survive in nutrient-limited environments^13,14^. *Astyanax mexicanus*, the Mexican cavefish, provides an intriguing example of successful adaptation to nutritional stressful environments^15^ (Figure 1A). Approximately 30,000-190,000 generations ago^1,16,17^, ancestral surface fish in the rivers of North-East Mexico were flooded into nearby caves. Faced with new selective pressures, including limited nutrient availability, these fish managed to survive and thrive in the caves^18^, evolving increased metabolic capacities^19–25^. Over time, they evolved into a cave-adapted ecotype known as blind cavefish while still maintaining reproductive compatibility with surface fish, making this species a powerful genetic model for studying evolutionary adaptation to an extreme environment. Additionally, certain genetic mutations were favored through natural selection^26^. Therefore, even under laboratory conditions, cavefish exhibit morphological, behavioral and physiological phenotypes that differ from surface fish, making them an ideal model organism to elucidate the genetic basis of extreme adaptations. To date, this species has been discovered in 35 caves in Mexico^27^. These adaptations occurred repeatedly in at least two independent events^1,28^ (Figure 1A), which also suggests potentially repeated adaptations of reproductive strategies. Understanding the reproductive biology of Mexican cavefish can provide valuable insights into the mechanisms that allow survival in extreme environments.

**Figure 1.**
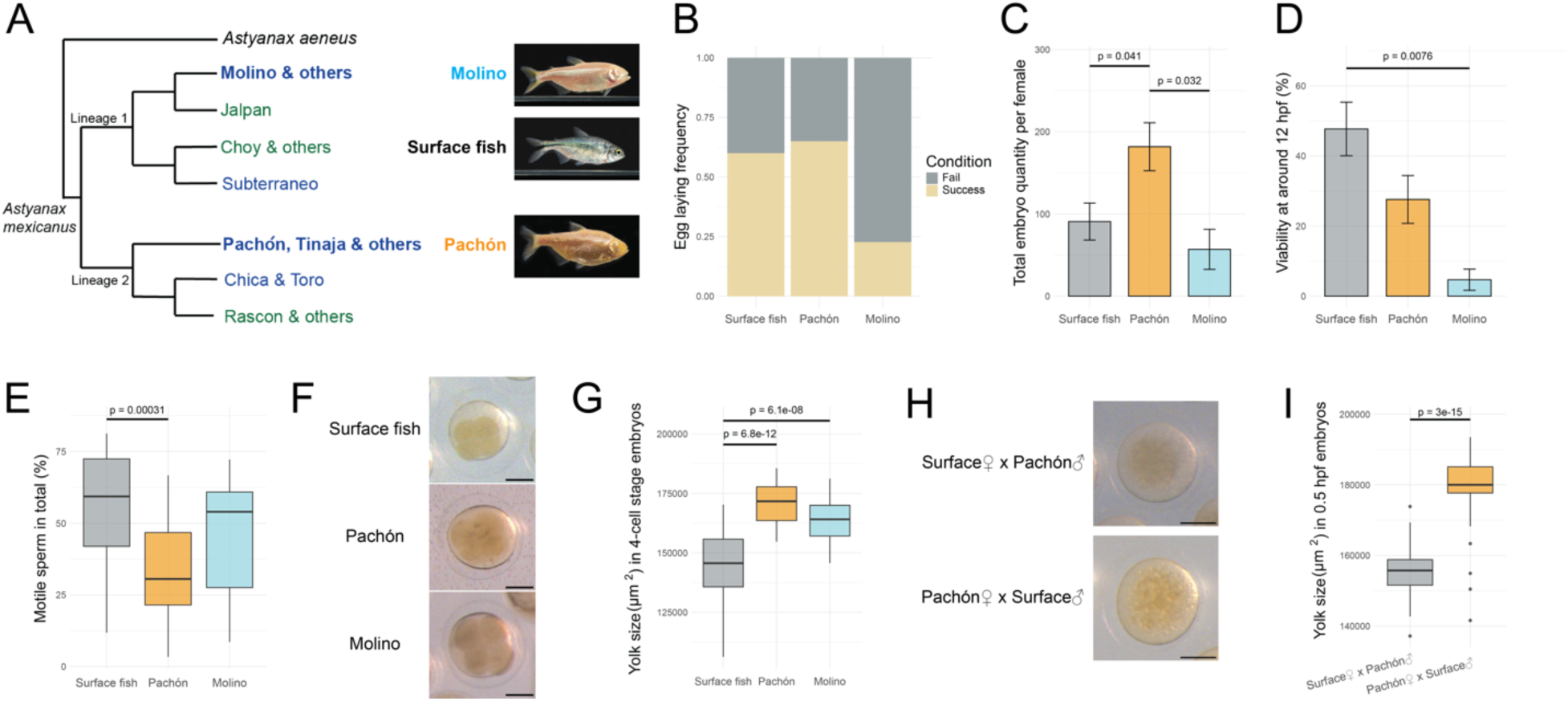
Independent cavefish populations have different reproductive output compared to surface fish. (A) Phylogenetic tree (left) was modified from Moran, *et al*^1^. Two independent evolutionary lineages of *A. mexicanus* are indicated. Green text denotes surface fish populations residing in the river, and dark blue text represents all cavefish populations. Representative images (right) from adult surface fish, Pachón and Molino show morphological differences among different populations. (B-D) The measurement of egg laying frequency, clutch size and embryo viability at the time of collection from the recordings of reproduction events in the lab. Egg laying frequency (B) was calculated from all spawning attempts (n=20, surface fish; n=20, Pachón; n=22, Molino). Clutch size (C) and embryo viability (D) were measured from successful spawning events (spawned eggs > 0; n=12, surface fish; n=13, Pachón; n=5, Molino). One way ANOVA was performed for statistical analysis for the comparisons of clutch size (F = 5.051, Pr(>F) = 0.0137). Pairwise comparisons were performed by Tukey’s HSD test. Kruskal-Wallis Test was performed for the comparisons of embryo viability (Kruskal-Wallis chi-squared = 9.4678, df = 2, p-value = 0.008792) among three populations. Pairwise comparisons were performed by Dunn’s test. P-values were adjusted with Bonferroni method. (E) Sperm motility of three fish populations measured during breeding cycles (n=28, surface fish; n=26, Pachón; n=16, Molino). One way ANOVA was performed for statistical analysis for the comparisons of clutch size (F = 8.659, Pr(>F) = 0.000452). Pairwise comparisons were performed by Tukey’s HSD test. (F) Representative images of 4-cell stage embryos (∼1.5 hpf) of three fish populations. Scale bar, 200 μm. (G) Yolk size measurements in 4-cell stage embryos of three fish populations (n=78, surface fish; n=18, Pachón and n=22, Molino). Measurements were only performed in orientations with all four cells on the top of the yolk. One way ANOVA was performed for statistical analysis for the comparisons among three populations (F = 41.14, Pr(>F) = 3.33e-14). Pairwise comparisons were performed by Tukey’s HSD test. (H) Representative images of Surface female x Pachón male hybrids and Pachón female x Surface male hybrids at 0.5 hpf. Scale bar, 200 μm. (I) Yolk size measurements in two hybrids (n=56 for Surface♀ x Pachón♂; n=49 for Pachón♀ x Surface♂). Mann-Whitney U Test was performed for statistically analysis for the comparisons between these two hybrids (W = 143, p-value = 3.002e-15).

Several studies have explored reproductive biology in *A. mexicanus*. Fieldwork has shown that cavefish can reproduce year-round in the wild^29^. Similarly, spawning and breeding events can be induced in the lab in surface fish, cavefish and between the two^30,31,32^. At the tissue level, primordial germ cell (PGC) migration, gonadal differentiation, and expression of certain sex-related genes have been characterized in Pachón cavefish using methods including PGC fluorescent labeling, hematoxylin and eosin (H&E) staining, and quantitative reverse transcription polymerase chain reaction (RT-qPCR)^33^. Surprisingly, many classical ovarian and testicular-specific genes were predominantly overexpressed in the testes^33,34^, indicating a potentially unique process of sex differentiation in *A. mexicanus* compared to some other teleost fish. Furthermore, light and electron microscopy imaging of sperm and oocytes provided the first cellular-level characterization of gametes in multiple surface and cave populations^35^. However, a more detailed comparison of reproductive output and characterization of gonads between surface fish and cavefish is still needed to better understand the evolution of reproductive biology and strategies in *A. mexicanus*, particularly how cavefish reproductive systems have adapted to extreme environments. These investigations could provide valuable insights into how environmental factors, such as nutrient availability, shape reproductive strategies in vertebrates.

To gain a more comprehensive understanding of reproductive biology in *A. mexicanus*, particularly focusing on female reproduction, we examined multiple aspects, including breeding output (number and composition of the eggs), as well as morphological and transcriptomic profiles in the ovary of surface fish and two independently derived cave populations, Pachón and Molino cavefish (Figure 1A). Finally, we explored whether and how these differences in the eggs or ovaries contribute to their reproductive abilities under nutrient-limited conditions. Through this comparative approach, we identified a few genes, including *insulin-like growth factor 1a receptor* (*igf1ra*), as potential candidates involved in starvation resistance in cavefish. Further, we found that during starvation, surface fish females failed to produce eggs, and exhibited a significant reduction in the number of mature oocytes. These changes were accompanied by the downregulation of *igf1ra* during starvation, which may lead to reduced steroidogenesis. In contrast, starved Pachón cavefish females were less affected by starvation. No significant gene expression changes were observed between control and starved Pachón cavefish, potentially indicating a protective mechanism that enhances starvation resilience in the cavefish reproductive system. Overall, this study offers a detailed examination of the *A. mexicanus* female reproductive system, shedding light on how female reproductive strategies adapt to environments with limited nutrient availability.

## Results

### Independent Cavefish Populations Have Different Reproductive Outputs Compared to Surface fish

Research on *Caenorhabditis elegans* and *Daphnia* have demonstrated that poor maternal nutritional environments lead to a reproductive trade-off, where mothers produce smaller clutch sizes but larger eggs^11,12,36^. This strategy is thought to help offspring better withstand nutritional stress by providing them with more resources per individual, thus potentially increasing their survival under adverse conditions. Based on this, we hypothesized that cavefish might exploit a similar strategy in response to an evolutionary adaptation to the nutrient-limited cave environment, even when housed in the laboratory. To assess the potential reproductive trade-off in *A. mexicanus*, we measured the number and size of eggs produced from different populations. We recorded female spawning frequency from all spawning attempts and measured clutch size and embryo viability from successful spawning events. Molino, but not Pachón cavefish, had less egg laying frequency compared to surface fish (Figure 1B). Clutch size was different in both cavefish populations compared to surface fish, with Pachón laying more eggs from a single brood whereas Molino had less compared to surface fish (Figure 1C). Additionally, a lower embryo viability was observed in both cavefish populations compared to surface fish at the time of collection (Figure 1D), which may be partially due to the low sperm mobility in cavefish (Figure 1E). These results suggest that Pachón and Molino cavefish have different but unique reproductive strategies compared to surface fish.

However, the results were counted and calculated from a limited number of broods (n=12, surface fish; n=13, Pachón; n=5, Molino), and could be affected by many individual variances. Therefore, we increased the number of replicates and estimated the number of eggs produced from each spawning attempt in our cavefish facility for one year (n=394, surface fish; n=291, Pachón; n=381, Molino). Combining one year of estimated recordings, we identified a similar trend of egg laying frequency, clutch size and embryo viability to support our earlier results (Supplementary Figure 1A-C). Notably, we observed some seasonal fluctuations in egg productions in Pachón and Molino cavefish populations with the most successful breeding peaks differing between populations (Supplementary Figure 1D), although only one successful egg laying event in Molino was recorded in March, which is insufficient to make a definitive statement. In contrast, age seemed not to affect egg production in cave populations but to some extent in surface fish (Supplementary Figure 1E).

Next, we used yolk size of early embryos as a proxy to compare the maternal investment in individual offspring among different populations. We found that 4-cell stage embryos (1 hour post fertilization (hpf)^37^) from both cavefish populations have larger yolk compared to surface fish (Figure 1F and 1G), and the difference extends to 14 hpf embryos (Supplementary Figure 1F and 1G). This observation aligns with a previous discovery from different surface and cave populations^38^. To exclude paternal influence, we measured the yolk size of surface fish and Pachón hybrids at 0.5 hpf. We found that hybrids produced from Pachón females exhibit larger yolk compared to the ones from surface fish females (Figure 1H and 1I).

Taken together, these results suggest that cavefish have a larger maternal deposition into individual offspring compared to surface fish, providing each embryo with more nutrient reserves. Aligning with our hypothesis, Molino produce smaller clutch sizes potentially to compensate for their larger egg sizes. Interestingly, Pachón exhibited both increased clutch size and egg size compared to surface fish, suggesting a generally higher maternal investment into reproduction.

### Egg Protein and Lipid Composition Indicate Morph Specific Evolutionary Adaptation in *A. mexicanus*

*Astyanax mexicanus* fish larvae start eating at 5.5 dpf^37^. Prior to this, the early embryos, like in many other oviparous teleost fish, rely on the yolk provided by the mother as their sole nutrient source for sustenance^39^. Therefore, yolk composition is essential for later-life development. Previous studies have shown that overall chemical compositions of eggs including protein and fat percentage is similar between surface fish and Pachón cavefish^38^. However, when examining specific yolk components, transcriptomic comparisons between 2-cell stage embryos from surface fish and Pachón cavefish revealed some differences in maternal RNA that are associated with signaling and cell interactions^40^. While transcriptomic data provides insights into maternal RNA deposition, it does not capture the full spectrum of yolk components. Many proteins, lipids, and small molecules, instead of being synthesized *de novo* within the eggs, are produced in other tissues or somatic cells in the ovary before being deposited into the eggs^41^. Dynamics of these components may not necessarily correspond with differences at the mRNA level. Therefore, to expand on the understanding of yolk composition, we used untargeted proteomics and lipidomics approaches to analyze protein and lipid profiles of maternal deposit in surface fish and cavefish using 2-cell stage embryos (Figure 2A), based on the assumption that embryonic proteomic profiles may not vary too much within the first hour post-fertilization^42^. Protein samples were labeled with Tandem Mass Tag (TMT) system^43^ for accurate protein quantification and comparisons among samples. Lipid samples were analyzed in both positive and negative ion mode. After quality control, a total of 1446 proteins, 2819 lipid species and related chemicals were detected. A pairwise differential expression analysis among three populations^44,45^ was performed using median-normalized counts.

**Figure 2.**
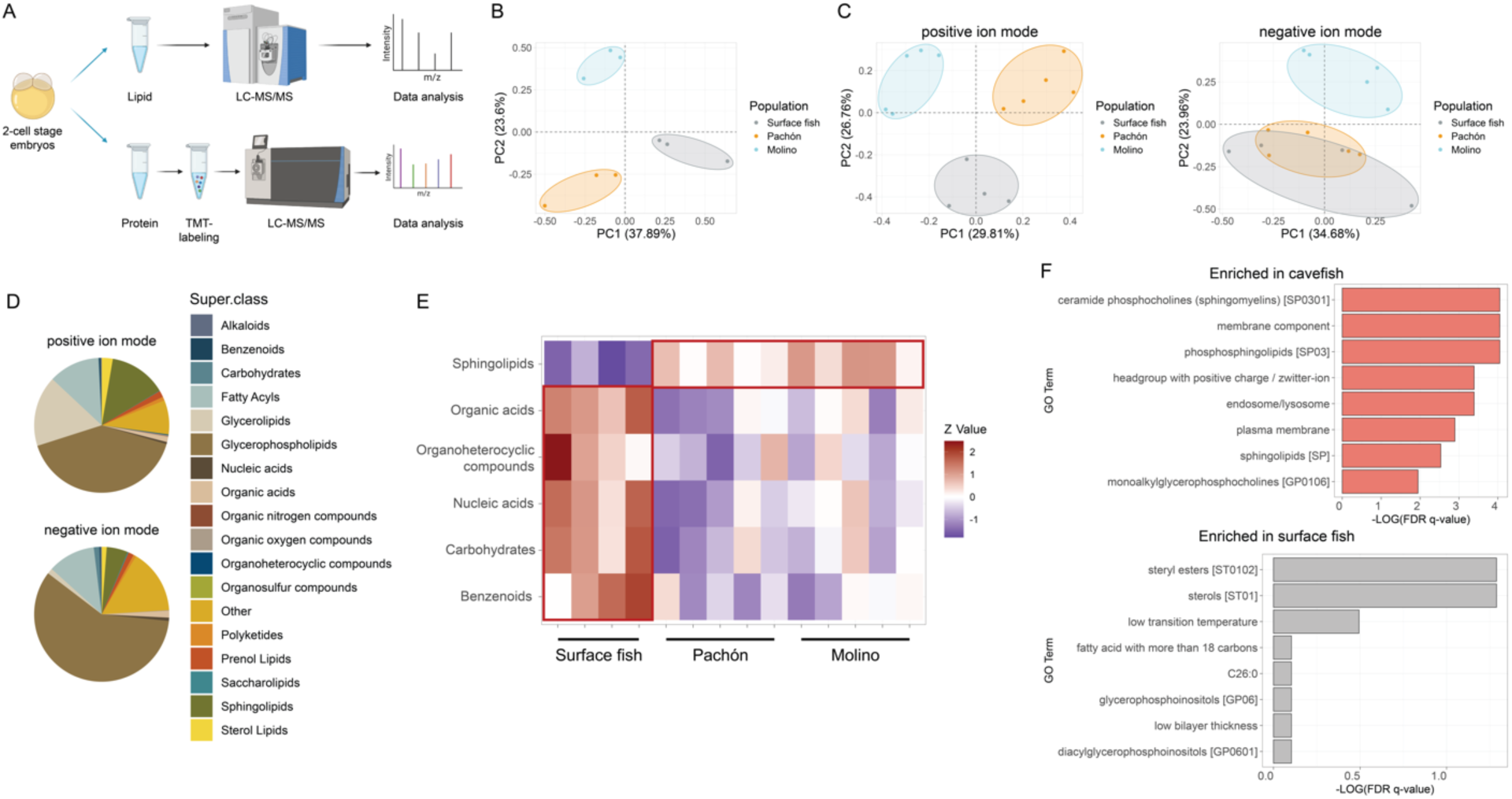
Unique yolk protein and lipid profiles between surface fish and cavefish. (A) Experimental schematics for protein and lipid extraction from 2-cell stage embryos. (B) PCA plot of proteomics results. (C) PCA plot of lipidomics results in positive and negative ion mode. (D) Pie plot of lipid super classes in 2-cell stage embryos of three populations in *A. mexicanus* in positive and negative ion mode. (E) Enriched lipid super classes in surface fish or cavefish. Benzenoids abundance was measured from negative ion mode. The abundance of other five super classes were measured from positive ion mode. (F) Lipid ontology analysis of individual lipid species with higher or lower abundance in cavefish compared to surface fish.

Principal component analysis (PCA) from protein samples of *A. mexicanus* 2-cell stage embryos showed a clear separation between fish morphotypes (surface fish / cavefish) but also between independently derived cavefish populations (Figure 2B). A total of 111 overlapping proteins were found to be commonly increased (65) or reduced (46) in both cavefish populations compared to surface fish (Supplementary Table 1, Supplementary Figure 2A, highlighted in red, log2FC ≥ 0.5 and padj ≤ 0.05). Notably, several candidates were found to exhibit the same pattern as in the mRNA level^40^. For example, compared to surface fish, cavefish 2-cell stage embryos have reduced protein and mRNA levels of metabolic genes such as malate dehydrogenase 1Aa, NAD (Mdh1aa), IMP (inosine 5,-monophosphate) dehydrogenase 1b (Impdh1b) and Isocitrate dehydrogenase (NADP(+)) 1 (Idh1), while increased protein and mRNA levels of several peptidases and hydrolases including asparaginase and isoaspartyl peptidase 1 (Asrgl1), sterile alpha and TIR motif containing 1 (Sarm1) and peptidase, mitochondrial processing subunit alpha (Pmpca). Aside from the examples listed above, most other candidates identified in our study were either not differentially expressed at the mRNA level or displayed opposite trends between protein and mRNA levels. These highlighted that our proteomics approach captured some novel protein candidates that were not identified from transcriptomic analysis. A full list of proteins with commonly increased or reduced abundances in both cavefish populations can be found in supplementary table 1.

The major yolk precursor protein, vitellogenin (Vtg), is an example with low mRNA levels yet high protein abundances in the yolk. Vtgs are primarily synthesized in the liver, transported to the ovary through the bloodstream, absorbed by oocytes via endocytosis, and subsequently incorporated into the yolk as major component^41^. Our proteomics analysis revealed that the abundance of Vtgs accounts for 30% of the total identified proteins, and their high abundance may have hindered the detection of low-abundance proteins. Vtgs are the primary source of amino acids for developing embryos^41^. Additionally, Vtgs have been shown to carry lipids and vitamins into the yolk and even act as immune effectors^46,47^. Given their function in metabolism and transgenerational immunity, Vtgs were selected as our initial targets for proteomic analysis. In many oviparous animals, such as teleost fishes, reptiles, and chickens, at least two types of Vtgs have been identified with different combinations of protein domains^48^. In *A. mexicanus*, several Vtg proteins exhibited significantly higher or lower abundance in cavefish compared to surface fish. Specifically, we observed increased levels of Vtgs with NCBI protein ID KAG9266789.1 and KAG9266788.1, and reduced levels of Vtg with ID XP_022535137.2 in cavefish compared to surface fish (Supplementary Table 1). However, Vtg types have not been fully categorized in *A. mexicanus*, and improved proteome annotation would provide better insight into abundance differences and potential functional differences of specific yolk protein variants between surface fish and cavefish.

Next, we focused on the lipid composition of the yolk using lipidomics. Over three-quarters of all detected lipid species were identified in positive ion mode. PCA plots of lipid profiles from positive ion mode showed clear clustering for each fish population (Figure 2C, left). In contrast, surface fish and Pachón cavefish lipid profiles obtained from negative ion mode were very similar to each other (Figure 2C, right), while Molino cavefish clustered apart. Using a combination of annotation tool RefMet^49^, manual annotation from LIPID MAPS^50–52^ (https://www.lipidmaps.org) and PubChem^53^ (https://pubchem.ncbi.nlm.nih.gov), we classified all identified lipid species and related chemicals into 18 super classes (Figure 2D). Our untargeted lipidomic approach found that the largest components in *A. mexicanus* embryos were glycerophospholipids, rather than sterol lipids found in zebrafish embryos^54^. These differences may be attributed to species-specific variations between zebrafish and *A. mexicanus*, biases between targeted and untargeted lipidomic approaches, or technical differences in sample preparation. For example, we did not remove the chorion prior sample preparation, which might have increased the proportion of membrane lipids, such as glycerophospholipids (while the zebrafish study did not specify whether the chorion was included or not).

To examine the differences in lipid compositions between surface fish and cavefish, we first compared the lipid abundance across each super class. We found that cavefish exhibited significantly higher levels of sphingolipids, but lower levels of benzenoids, carbohydrates, nucleic acids, organoheterocyclic compounds and organic acids compared to surface fish (Figure 2E). To identify specific lipid species contributing to these differences, we performed pairwise comparisons between surface fish and cavefish populations. Lipid species or related chemicals that are commonly enriched in surface fish or both cavefish were identified (Positive ion mode: Supplementary Figure 2B; negative ion mode: Supplementary Figure 2C, common genes highlighted in red). Lipids with differential abundances between surface fish and cavefish in positive ion mode were categorized by super classes (Supplementary Figure 2D). The ratio of sphingolipids increased in lipid profiles enriched in cavefish, while the ratio of fatty acyls, nucleic acids, organic acids and sterol lipids increased in lipid profiles enriched in surface fish, consistent with the findings in Figure 2E.

To explore the function of enriched lipids in either surface fish or cavefish, we conducted lipid ontology enrichment analysis through the web-based tool LION^55^ (Figure 2F). We found that sphingomyelins (SM) were enriched in cavefish compared to surface fish (Figure 2F, top). While primarily known as membrane structural lipids^56^, SM and other sphingolipids play roles in various cellular signaling pathways, including those regulating lipid metabolism^57^, stress and immune responses^58,59^. The higher maternal provision of SM and other sphingolipids in cavefish might underlie key evolutionary adaptations to cave environments with different microbiome and parasite assemblages^27,60,61^ and reduced nutrient availability. Additionally, although not statistically significant, steryl esters were found to be enriched in surface fish compared to cavefish (FDR q-value = 0.0515, Figure 2F, bottom). A previous metabolomics study in the brains, livers, and muscles of adult *A.mexicanus* reported that cholesteryl esters were less abundant in cavefish than surface fish under certain feeding conditions^24^. Thus, our analysis provides another evidence for this unexpected finding of less active steroid metabolism in cavefish than surface fish, though the underlying mechanisms and functions need further investigation.

Altogether, we characterized the protein and lipid profiles in the eggs of three populations of *A. mexicanus*, providing additional information about chemical compositions of the yolk in each population. We identified distinct yolk protein and lipid profiles between surface fish and cavefish, which may confer adaptive advantages for living in different natural environments.

### Oocyte Morphology Across Developmental Stages are Similar Between Surface fish and Cavefish

Yolk components are synthesized within oocytes and absorbed from the plasma at different stages of oocyte maturation^41,62^. The efficiency of synthesis, uptake and processing of nutrients will affect their abundance in the yolk and subsequently influence offspring fitness^63^. To explore potential functional differences during female fish oocyte development that contribute to the observed variations in maternal provisioning between surface fish and cavefish, we first examined the ovary morphology from three populations of *A. mexicanus*.

In zebrafish, oocytes progress through different stages of oogenesis to prepare for ovulation^64^. For *A. mexicanus*, we followed a similar classification system using zebrafish oocyte staging categories^64^. Previous research has shown the morphological and molecular characteristics of early oogenesis in Pachón cavefish, from PGCs to early stage I oocytes (primary growth stage) during PGC migration^33^. To expand upon these findings by characterizing later oocyte developmental stages, we performed H&E staining on paraffin sections of age-matched adult fish ovaries (Figure 3A). This allowed us to capture stage Ib (follicle phase of primary growth), stage II (cortical alveolus stage) and stage III (vitellogenesis) and later stages of oocytes, along with surrounding somatic cells. We trained a YOLOv8 object detector^65^ to find oocytes belonging to one of three stages from H&E stained images. This was applied to thousands of images, where the sizes were quantified (Figure 3B). While acknowledging the inherent technical limitations of this approach, such as the fact that oocytes are not perfectly round in tissue sections and that we did not always capture the largest cross-sections, our estimate provides valuable insights. Overall, we did not observe substantial differences in oocyte sizes among populations except for a slightly larger mature oocyte size in cavefish compared to surface fish.

**Figure 3.**
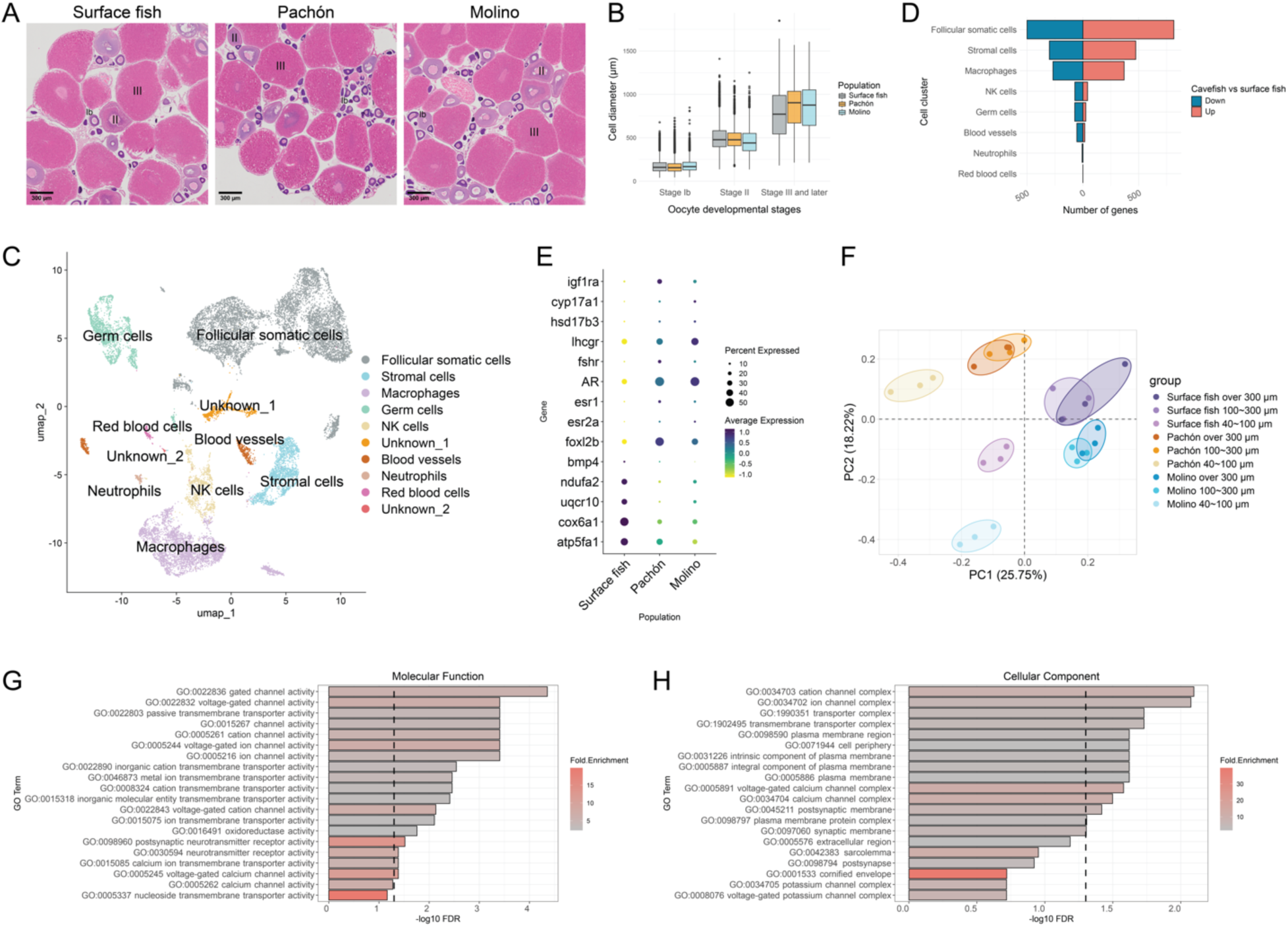
Ovary characterization revealed unique characteristics in cavefish compared to surface fish. (A) Representative H&E-stained ovary sections from three fish populations at 500 to 600 dpf. Scale bar, 300 μm. Different stages of oocytes (stage Ib, II and III) are labeled. (B) Oocyte size (diameter) measurement in different developmental stages of surface fish, Pachón and Molino. (C) Single-cell UMAP plot of the adult fish ovaries from *A. mexicanus*. Cells are color-coded by computationally determined cell clusters. (D) DEGs between surface fish and cavefish in eight annotated cell clusters. (E) Dot plot of gene expression in follicular somatic cells. Selected genes are involved in hormone responses, synthesis or OXPHOS pathways. (F) PCA plot of transcriptomic profiles of oocytes at different size categories in three fish populations. (G) Molecular function and (H) Cellular component GO term analysis of commonly upregulated genes in oocytes larger than 300 µm of both cavefish compared to surface fish. The dash line indicates significant threshold (FDR = 0.05).

### Ovary Expression Profiles Reveals Cavefish-specific Reproductive Adaptation

To further investigate whether there are molecular differences between surface fish and cavefish ovary, we separated ovarian cells into different size ranges. We performed single cell RNA sequencing (scRNA-seq) on cells with a diameter smaller than 40 μm due to technical limitations to capture all somatic cell types in the ovary and early germ cells, and bulk RNA sequencing (RNA-seq) on cells larger than 40 μm to obtain a more comprehensive overview of gene expression profiles from larger oocytes.

Cells smaller than 40 μm were clustered into ten distinct populations using uniform manifold approximation and projection (UMAP; Figure 3C). We assigned cell types to different clusters based on known markers for different cell types, namely, *deadbox helicase 4* (*ddx4*, or *vasa*) for germ cells^66^; *gonadal soma derived factor* (*gsdf*)^67^, *hydroxysteroid (17-beta) dehydrogenase 1* (*hsd17b1*)^68^, *cytochrome P450, family 19, subfamily A, polypeptide 1a* (*cyp19a1a*), *hydroxysteroid (17-beta) dehydrogenase 3* (*hsd17b3*)^68^, *luteinizing hormone/choriogonadotropin receptor* (*lhcgr* or *lhr*)^69^ and *cytochrome P450, family 17, subfamily A, polypeptide 1* (*cyp17a1*)^70^ for early to late stages of follicular somatic cells; *collagen, type I, alpha 1a* (*col1a1a*) for stromal cells^71^; *friend leukemia integration 1 transcription factor* (*fli1*) for blood vessels^72^; *band 3 anion exchange protein-like* (*slc4a1a*) for red blood cells^73^; *macrophage expressed 1* (*mpeg1*) for macrophages; *myeloid-specific peroxidase* (*mpx*) for neutrophils^74^ and *interleukin 2 receptor, beta* (*il2rb*) for NK cells^75^ (Supplementary Figure 3A). Similar cell types were found among zebrafish^71^ and three populations in *A. mexicanus* ovaries (Supplementary Figure 3B). Cell proportion analysis using scCustomize^76^ and scProportionTest^77^ suggested that both Pachón and Molino cavefish exhibited more follicular somatic cells and blood vessels, while less immune cells compared to surface fish (Supplementary Figure 3C-E). To identify which cell cluster contributes most to the differences between surface fish and cavefish ovaries, we conducted differential gene expression analysis across all eight annotated cell clusters. A large number of genes were commonly differentially expressed between surface fish and both cavefish populations in follicular somatic cells (Figure 3D). Coupled with its higher proportion in cavefish compared to surface fish, these results suggest that follicular somatic cells may play an important role in cavefish-specific reproductive adaptation. Therefore, we focused on follicular somatic cell population to explore potential implications of the differences between surface fish and cavefish reproductive strategies.

Follicular somatic cells, composed of follicle cells and theca cells surrounding oocytes, provide support for oocyte development and maturation. In teleost fish, follicle cells are homologous to granulosa cells in mammals, forming the inner follicular layer, while theca cells form the outer layer, separated by a basement membrane^64^. In mammals, these follicular somatic cells play crucial roles in hormone production, such as estrogen synthesis^78^, and provide nutrients to support the developing oocytes^79^. Recent scRNA-seq analysis in zebrafish ovary suggested that some of these functions may be conserved in fish^71^.

To explore the differences in follicular somatic cells between surface fish and cavefish, we first subclustered these cells and performed a trajectory analysis using pre-follicle cells marked by *LIM homeobox 9* (*lhx9*)^71^ as starting population (Supplementary Figure 3F). This allowed us to capture follicular somatic cells from early to late stages across all populations, while no obvious compositional variation in cell types was observed. Therefore, we further examined molecular-level differences. Specifically, we performed gene ontology (GO) enrichment analysis using a web-based tool^80,81^ on 814 commonly upregulated genes and 500 commonly downregulated genes in follicular somatic cells of Pachón and Molino compared to surface fish (Supplementary Figure 3G and 3H). Cavefish follicular somatic cells showed upregulation in genes enriched in pathways related to cellular response to hormone stimulus, including *cyp17a1*, *hsd17b3*, *lhcgr*, *follicle stimulating hormone receptor* (*fshr*), *androgen receptor* (*ar*), *estrogen receptor* (*esr1*), *estrogen receptor beta-1-like* (*esr2a*) and *forkhead box L2b* (*foxl2b*) (Figure 3E). Meanwhile, *bone morphogenetic protein 4* (*bmp4*), which suppresses androgen production^82^ and participates in the TGF-beta signaling pathway, was downregulated in cavefish compared to surface fish (Figure 3E). These findings suggest that cavefish have an enhanced capacity for hormone regulation compared to surface fish. Additionally, cavefish follicular somatic cells exhibited reduced expression of genes involved in oxidative phosphorylation (OXPHOS) pathway compared to surface fish, including those encoding components of mitochondrial complexes such as NADH dehydrogenase, cytochrome c reductase, cytochrome c oxidase and F-type ATPase (Figure 3E), suggesting that cavefish may have a lower energy demand than surface fish into reproductive processes. Overall, these findings highlight distinct energy production and hormone regulation strategies in surface fish and cavefish follicular somatic cells, potentially contributing to reproductive success in their respective environments.

Apart from somatic cells, different stages of oocytes were isolated from the right-side ovary of the same fish used for scRNA-seq. Bulk RNA-seq was performed to assess transcriptomic profiles of the oocytes larger than 40 µm. More specifically, we separated cells into four categories based on size: smaller than 40 µm; 40∼100 μm; 100∼300 μm and larger than 300 μm. According to our scRNA-seq data, cells smaller than 40 μm were mainly somatic cells, with less than 20% identified as germ cells (Supplementary Figure 3C). Based on approximate size characterization (Figure 3B), cells between 40 and 100 μm largely consisted of stage I oocytes, while those between 100 and 300 μm were a mix of stage I and mostly stage II oocytes. Cells larger than 300 μm predominantly contained stage III oocytes and later-stage oocytes undergoing vitellogenesis, though some stage II oocytes were also present. PCA revealed a clear separation of transcriptomic profiles between cells smaller and larger than 40 µm (Supplementary Figure 4A), consistent with previous findings in zebrafish^83^, likely reflecting the difference between somatic cells and oocytes. We further conducted PCA specifically on oocytes and found that each fish population clustered separately (Figure 3F). Notably, Pachón and Molino cavefish oocytes clustered in opposite directions from surface fish samples. In contrast, proteomics data showed that samples from the cavefish populations clustered together (Figure 2B). This discrepancy between the transcriptomic and proteomic profiles may suggest either inherent differences in the two data types or, more intriguingly, that the distinct transcriptomic profiles of Pachón and Molino cavefish lead to converging proteomic outputs and ultimately similar cave phenotypes. Additionally, oocytes between 100 and 300 μm clustered together with those larger than 300 μm, indicating similar transcriptomic profiles throughout vitellogenesis.

To investigate cave-specific differences at different oogenesis stages, we conducted GO term analysis on common differentially expressed genes (DEGs) between surface fish and Pachón or Molino across different oocyte categories. In oocytes larger than 300 μm, we observed that upregulated genes in both cavefish populations compared to surface fish were enriched in molecular functions related to cation channel activity (Figure 3G) and in cellular components such as cation channel complex and post synaptic membrane (Figure 3H). Similar pathways were also identified in oocytes between 100 and 300 μm (Supplementary Figure 4B). Fish ovaries can be innervated and regulated by neuronal signals^84^. These findings suggested that stage II and later-stage oocytes in cavefish may exhibit more neuronal connections or innervations compared to surface fish. However, our scRNA-seq analysis did not reveal a neuronal cell population, nor did we find evidence of dopaminergic neurons near the oocytes using tyrosine hydroxylase as the marker (data not shown). This suggests that the synapse-related genes may serve alternative intra- or intercellular communication functions. A previous study in bovine ovaries highlighted the role of synapse-like structures of transzonal projections in small molecules transfer such as RNA between cumulus cells and oocytes^85^. Based on this, we hypothesized that cavefish may exhibit enhanced nutrient transfer efficiency between oocytes and their surrounding microenvironment, which may contribute to the observed reproductive advantages in cavefish egg size (Figure 1F and 1G).

To investigate this hypothesis, we performed electron microscopy imaging around the vitelline envelope (VE) (or zona pellucida) regions at different stages of oogenesis (Supplementary Figure 4C-K), to investigate potential differences in intercellular communication between follicle cells and oocytes. We observed the microvilli from both follicle cells and oocytes. These microvilli were seen extending into the surface of the oocytes in stage III or later oocytes (500∼600 μm) in all three populations (Supplementary Figures 4E, 4H and 4K), a feature not observed in zebrafish^64^. However, there were no significant morphological differences among the three populations of *A. mexicanus* in terms of pore canal or microvilli numbers. This suggests that the higher nutrient reserves in cavefish eggs may not be related to pore canal structures, and other mechanisms contributing to this phenomenon require further exploration.

Another possible mechanism involves the increased expression of cation channels, including Ca^2+^, K^+^, and Na^+^ channels, in cavefish compared to surface fish. Calcium homeostasis and oscillation in oocytes are well known to be critical for oocyte maturation and fertilization^86^. These ion channels traditionally play vital roles in cellular communication and signal transduction^87^. The increased expression of these channels may indicate a heightened level of ion exchange, facilitating a more active signal transduction network within the oocytes. This enhanced communication might also ensure proper intercellular communication between follicle cells and oocytes, potentially contributing to cave-specific reproductive strategies. Overall, we have generated a comprehensive database of transcriptomic profiles across various cell types/sizes in the ovary and different oocyte stages, which we hope will further the understanding of female reproduction in *A. mexicanus* and provide a valuable resource for future research.

### Cavefish Reproduction is Starvation Resistant

In the wild, cavefish can survive for long periods of time in nutrient-limited environments, yet still maintain the ability to reproduce year-round^29^, suggesting high starvation resilience in its reproductive system. To investigate how starvation affects the reproductive capabilities of surface fish and cavefish, we conducted a 2-month starvation experiment on age-matched females from surface fish and Pachón cavefish and induced spawning every month (Figure 4A). For clearer data visualization, we grouped results from starved fish at 1 and 2 months into “starvation group”, while results from non-starved fish and starved fish at 0 month (regular feeding, right before starvation started, see Figure 4A and Methods) were classified as the “control group”.

**Figure 4.**
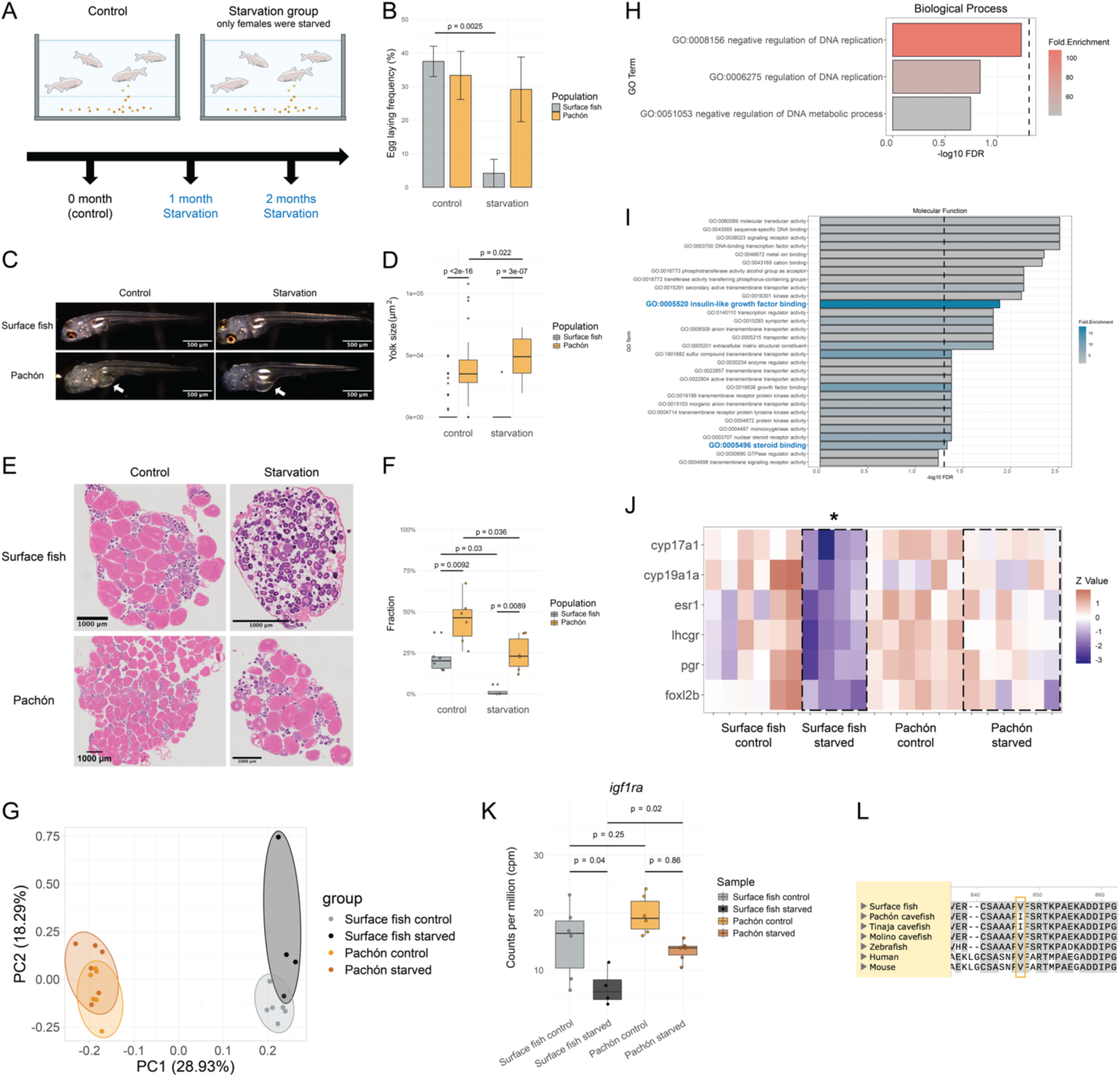
Pachón cavefish reproduction is starvation resistant. (A) Experimental set up for starvation and breeding experiment. Only female fish were starved. (B) Egg laying frequency of surface fish and Pachón under control and starvation. Control condition includes all time points from control groups and 0 month from starvation groups (n=12, surface fish control; n=12, Pachón control; n=6, surface fish starvation; n=6, Pachón starvation). Aligned Rank Transform (ART) ANOVA from ARTool v0.11.1^2^ in R 4.3.1 was performed for statistically analysis for the comparisons (Condition:population F = 7.4206, Pr(>F) = 0.0081883). Post-hoc tests were performed by built in art.con() function. (C) Representative lateral view of 6 dpf surface fish and Pachón offspring from control and starved fish. Scale bar, 500 μm. Arrows point to the yolk. (D) Yolk size measurement of 6 dpf surface fish and Pachón offspring from control and starved fish (up to 5 larvae were selected from the same spawning event; n=61, surface fish control; n=58, Pachón control; n=5, surface fish starvation; n = 25, Pachón starvation). Aligned Rank Transform (ART) ANOVA was performed for statistically analysis for the comparisons (Condition:population F = 1.7155, Pr(>F) = 0.19238). Post-hoc tests were performed by built in art.con() function. (E) Representative H&E-stained ovary sections from surface fish and Pachón under control and starvation. Scale bar, 1000 μm. (F) Fractions of stage III or later (vitellogenic) oocytes in surface fish and Pachón under control and starvation. Aligned Rank Transform (ART) ANOVA was performed for statistically analysis for the comparisons (Condition:population F = 0.0068952, Pr(>F) = 0.93473818). Post-hoc tests were performed by built in art.con() function. (G) PCA plot of transcriptomic profiles of ovary from surface fish and Pachón under control and starvation. (H) Biological Process GO term analysis of upregulated genes in starved surface fish ovary compared to surface fish control. (I) Molecular Function GO term analysis of downregulated genes in starved surface fish ovary compared to surface fish control. The dash line indicates significant threshold (FDR = 0.05). (J) Heatmap showing expression levels of genes that are hormone receptors or contribute to ovary maintenance across experimental groups. All genes shown in the figure were significantly downregulated in starved surface fish compared to surface fish control. (K) *igf1ra* expression level (cpm) across surface fish and Pachón experimental groups. Adjusted p-value was calculated from bulk RNA-seq analysis. (L) Amino acid alignment of *igf1ra* highlights the conserved V826I mutation in Pachón and Tinaja cavefish from *A. mexicanus*.

During 2 months of starvation, we observed that both starved surface fish and Pachón females experienced significant weight loss compared to their initial weight at the start of the experiment, whereas females in the control group gained weight. The weight losses were statistically significant in both surface fish and Pachón compared to their respective control groups (Supplementary Figure 5A). Males were only fasted for 5 days when introduced into the starved female tanks for the breeding cycle. Despite this short fasting period, surface fish males also exhibited weight loss compared to controls, though the reduction was less pronounced than in females (Supplementary Figure 5B).

As predicted by their natural environments, breeding and spawning in starved Pachón females were less affected by starvation compared to surface fish. Surface fish experienced a significant reduction in egg-laying frequency during starvation (p = 0.0025, ART ANOVA), whereas Pachón showed no such effect (Figure 4B). Only one starved surface fish female spawned once at 1 month after starvation. Clutch size and offspring survivability from approximately 12 hpf through 6 dpf were not significantly affected by starvation (Supplementary Figure 5C-E). Interestingly, we observed a significant increase in yolk size in 6 dpf offsprings from starved Pachón compared to their controls (Figure 4C and 4D), a phenomenon also reported in zebrafish upon starvation^9^. However, the body length of 6 dpf larvae was not significantly impacted by starvation (Supplementary Figure 5F). The enlarged yolk in 6 dpf larvae from starved females may indicate that starved Pachón laid bigger eggs than controls, or that offspring from starved females exhibit a slower yolk consumption rate. Either of these mechanisms would enhance offspring fitness in nutrient-limited environments. These findings suggested that Pachón reproductive ability may possesses a higher plasticity in adapting to nutritional stressful environments compared to surface fish.

Given these observations, important questions arise: How are Pachón cavefish able to lay eggs during starvation while surface fish are more impacted? Do the morphological and transcriptomic profiles of surface fish and Pachón ovaries change in response to nutrient deprivation? If so, what may contribute to their different responses? To uncover the potential mechanisms behind the starvation resilience in Pachón cavefish reproductive system, we aimed to explore both the morphological and transcriptomic profiles of the ovaries during starvation.

We first investigated morphological changes in the ovaries of surface fish and Pachón during starvation. At the end of the starvation period, we dissected the ovaries from all female fish and performed H&E staining on paraffin sections. The percentage of ovary weight to body weight was significantly higher in Pachón compared to surface fish under both feeding and starvation conditions (p = 0.0039 for control and p = 0.023 for starvation condition, Two-way ANOVA), suggesting an increased reproductive investment in Pachón relative to surface fish (Supplementary Figure 5G). Additionally, ovary weight percentage decreased under starvation in both morphs (Supplementary Figure 5G). Interestingly, we observed an opposite correlation in fat and ovary weight percentage between surface fish and Pachón, where surface fish showed a positive correlation while Pachón displayed a negative correlation (Supplementary Figure 5H and 5I). This suggests different energy allocation strategies between self-maintenance and reproduction in surface fish and Pachón. Upon dissection, most ovaries from starved surface fish displayed a very transparent appearance, with almost no visible opaque eggs, suggesting immature eggs. Consistent with these observations, no mature or vitellogenic oocytes were found in starved surface fish (Figure 4E), except in one individual that was able to spawn during the experiment (Figure 4F). The reduction of mature oocytes likely explains why most surface fish were unable to spawn during starvation.

### Potential Role of IGF Signaling in Mitigating Starvation Effects on Cavefish Reproduction

We next performed bulk RNA-seq of the ovaries from all experimental female fish and conducted PCA (Figure 4G). Samples from control and starved Pachón clustered closely together, suggesting similar transcriptomic profiles between these conditions. This aligns with our findings that reproductive ability in Pachón were not significantly affected by starvation. Conversely, starved surface fish samples showed large variation among replicates and separated from surface fish under control conditions. Notably, the one starved surface fish sample that clustered with control samples came from the female that successfully spawned once during starvation. Differential gene expression analysis did not reveal any significantly DEGs (adjusted p-value < 0.05) between control and starved Pachón ovaries. We then conducted GO analysis on DEGs between control and starved surface fish. Although not statistically significant, upregulated genes in starved surface fish were enriched in biological process related to negative regulation of DNA replication (Figure 4H), suggesting potential decreased cell proliferation in surface fish during starvation. Downregulated genes in starved surface fish were enriched in multiple molecular functions, including insulin-like growth factor binding, nuclear steroid receptor activity and steroid binding (Figure 4I). More specifically, we found a significant decrease of genes related to ovarian development in surface fish during starvation, whereas this pattern was absent in Pachón (Figure 4J). Some of these genes were also upregulated in cavefish follicular somatic cells compared to surface fish (Figure 3E), with *igf1ra* being one of the shared genes across both comparisons.

*igf1ra* is predominantly expressed in somatic support cells, including follicular somatic cells and stromal cells (Supplementary Figure 5J), and shows higher expression in cavefish follicular somatic cells compared to surface fish (Figure 3E). We observed significant downregulation of *igf1ra* in surface fish during starvation (p = 0.04), while no such reduction was seen in Pachón (p = 0.86) (Figure 4K). This starvation response was also observed in another independent starvation experiment with a separate batch of surface fish and Tinaja cavefish, an additional cave population (Tinaja cavefish evolutionary position is shown in Figure 1A, Supplementary Figure 5K, p = 0.071 in surface fish). Insulin-like growth factor 1 (IGF-1) production is sensitive to nutritional environment^88^. Insulin-like growth factor 1 receptor (IGF1R) protein level in the growth plate was reduced during food restriction^89^, consistent with the significant reduction in surface fish fertility upon starvation. Intriguingly, a previous study has shown that mice with granulosa cell-specific knockout IGF1R failed to ovulate, which was accompanied with the downregulation of genes involved in steroidogenesis such as *LHCGR* (or *LHR*) and *Cyp19a1* (or aromatase)^90^. The phenotype and molecular changes we observed in starved surface fish closely resemble those seen in the IGF1R knock out mice, including impaired egg-laying ability (Figure 4B) and downregulation of *lhcgr* and *cyp19a1a* (Figure 4J). Moreover, we identified a coding variation (V to I) in the exon of *igf1ra* in Pachón and Tinaja cavefish (Figure 4L), which could potentially alter its function. These findings suggest that the upregulation of *igf1ra* or IGF signaling in cavefish follicular somatic cells or its function may provide a buffering mechanism, allowing cavefish to better withstand starvation. As a result, other genes that are expressed in follicular somatic cells and essential for female reproduction, such as *cyp19a1a*, may be less affected by nutritional stress.

## Discussion

In this study, we investigated reproductive biology in the Mexican cavefish focusing on spawning and breeding output, as well as morphological and transcriptomic profiles of ovaries. We also explored the mechanisms underlying starvation resilience in cavefish females. Our findings revealed that Pachón and Molino produced different number of eggs compared to surface fish. Eggs from both cave populations were larger than surface fish with unique maternal deposition. Additionally, our comprehensive transcriptomic analysis identified DEGs across different cell types between surface fish and cavefish ovaries, highlighting candidate genes involved in hormone regulation or ion channels that may provide a buffering mechanism in cavefish reproductive systems. These mechanisms likely enable cavefish to withstand long periods of starvation while continuing to reproduce, in contrast to surface fish, which cannot maintain reproductive activity under the same conditions.

### Quantity and quality of eggs – a trade off in reproductive strategy

A tradeoff in reproductive strategy between quantity and quality of offspring was proposed by the r/k selection theory^91^. According to this theory, species under r-selection (reproduction) prioritize producing a large number of offspring, with relatively less parental care. This strategy provides adaptive advantages in unstable environments. In contrast, K-selection (carrying capacity) refers to a greater investment in fewer offspring, enhancing competitive ability in more predictable environments^92,93^. The life histories of many animals are good examples of conventional r/k selection^94^, while the relationships among life-history traits can be more complex in others^95^. Compared to the surface rivers, caves have lower species diversity^96^ and more stable abiotic conditions, such as temperature^97^. Therefore, we hypothesized that cavefish, living in a more stable environment, might exhibit traits of K-selection, while surface fish show traits of r-selection.

Since the Mexican cavefish is an oviparous fish species, we measured the number and the size of the eggs as proxies for accessing the number of offspring and maternal investment into each offspring. Over the course of one year breeding recordings in the lab, we found that Pachón produced a larger clutch size and larger eggs than surface fish, whereas Molino have a smaller clutch size but larger eggs compared to surface fish. The larger egg size in cavefish aligns with previous findings in several other cave populations^37,38^, and supports our hypothesis of K-selected traits in cavefish. However, our recorded clutch size differences between surface fish and Pachón contradicts a previous finding^32^. One possible explanation for this discrepancy could be the use of different surface fish populations for comparisons. Instead of surface fish originating from San Solomon Spring, Balmorhea State Park, Texas, USA, our study used Río Choy population originally from Mexico. Indeed, both studies reported similar egg counts for Pachón, with an average of 1442 eggs per 7 females in their findings^32^, compared to an average of 182 eggs per female in our findings shown in Figure 1C. These suggested that clutch size and egg production may differ significantly across distinct surface fish populations, as expected given local adaptation. The unique local environments of different caves may have led Pachón and Molino cavefish to adopt distinct reproductive strategies. Moreover, it is important to recognize that laboratory conditions differ significantly from the wild. With different nutritional environments and interspecific interactions in the wild, all three populations may reproduce differently. However, our finding provided valuable insight into how these three populations reproduce under standardized and controlled conditions. Overall, our findings supported the notion that Pachón and Molino females invest more into their offspring, as evidenced by larger egg sizes in cavefish or hybrids with cavefish mothers (Figure 1F-I). Under sufficient nutrient availability, Pachón also appear to invest more into offspring quantity, while Molino reduce egg number, potentially balancing reproductive energy expenditure.

The cave environment is nutrient-limited compared to river habitats. Our initial hypothesis was that cavefish might breed only during the rainy season, when food could potentially be carried into the caves by floods^18^. However, evidence suggested that cavefish can breed year-round^29^, indicating that cavefish is capable of reproducing despite nutrient scarcity. Comparative analysis of female reproductive abilities under starvation conditions between surface fish and Pachón support the idea that cavefish exhibit a stronger resilience to reproduce in a low nutrient condition (Figure 4B).

This raises intriguing question: For females that spawned during starvation, were the mature oocytes present in the ovary before starvation began, or were they produced during the starvation period? Since spawning was induced prior to starvation (Figure 4A and Methods), we assume that only a limited number of mature oocytes were present in the ovaries at the onset of food deprivation. Therefore, most mature eggs laid by starved females or observed in the ovaries were likely produced and developed during starvation period. However, it is possible that some mature oocytes were retained after the initial induced spawning, did not degenerate or reabsorbed during starvation, and were subsequently ovulated and recorded during starvation period. Future studies are needed to further characterize the timeline of oogenesis and its response to starvation in *A. mexicanus*.

We identified one candidate gene, *igf1ra*, which is highly expressed in cavefish follicular somatic cells compared to surface fish (Figure 3E) and significantly reduced in surface fish during starvation (Figure 4K and Supplementary 5K). Considering IGF1R sensitivity to nutritional environments and its regulatory role in reproduction, these findings indicate that the increased expression or coding variation within *igf1ra* may contribute to the starvation resilience observed in cavefish, however functional validation in future studies are needed.

In this study, we mainly focused on *igf1ra* as a candidate gene of interest. The differences of its gene expression level changes between surface fish and Pachón during starvation prompted us to consider the possibility of selected mutations in upstream regulators or epigenetic modifications of *igf1ra*. It is also likely that this starvation protection is not limited to the ovary, but is widespread across various tissues. In adaptation to prolonged food scarcity in the wild, cavefish exhibit distinct metabolic activities, which may offer advantages to survive under starvation conditions^98^. These metabolic adaptations likely existed across multiple tissues. The synthesis and circulation of hormones or other proteins that are essential for ovary development and maintenance^7,41,84^, across organs like brain, liver and adipose tissue, may also be affected by starvation. Therefore, starvation resilience in other organs may provide a buffering microenvironment in cavefish, allowing the ovaries to be less sensitive to nutritional challenges at the body level.

Finally, another compelling observation was the single instance of a surface fish female that successfully spawned once after one month of starvation. We hypothesized that surface fish, as a population, may have a high plasticity in response to environmental changes, which could play a role in its rapid, unique and repeated cave adaptation.

## Materials and methods

### Fish husbandry

The photoperiod in the fish room and the embryo incubator followed a 14:10 light-dark cycle. The embryo incubator was set to 23°C. Fish older than 3 dpf were housed on Pentair recirculating racks equipped with biological and mechanical filtration, as well as UV disinfection. The fish stocks that were not used for breeding, were kept in polycarbonate 10L tanks at a stocking density of around 15 adult fish per tank. The ideal water quality setpoints were a temperature of 23°C, specific conductance of 800 µS, pH of 7.65, with total ammonia nitrogen, nitrite, and nitrate all at 0 mg/L, and dissolved oxygen levels above 90%. Standard fish were fed twice a day. The first feed consisted of Mysis (96908, Hikari USA), and the second feed was Gemma Silk 0.8 mm (Skretting).

Breeder fish were kept in 38L glass tanks with a maximum stocking density of 50 cavefish or 35 surface fish per tank. They were fed three times a day from Monday to Friday if not undergoing a breeding cycle. The first feed was Mysis, followed by two feeds of Gemma Silk 0.8 mm. On Saturday and Sunday, non-breeding fish received one feed of Mysis. During a breeding week, breeder fish were fed Gemma Silk 0.8 mm once a day from Monday to Friday, and on Saturday and Sunday, they received one feed of Mysis and two feeds of Gemma Silk 0.8 mm. All animal husbandry methods were approved by the Institutional Animal Care and Use Committee (IACUC) of the Stowers Institute for Medical Research under protocol 2021-122.

### Breeding

In our laboratory conditions, the breeding events are triggered by mimicking the temperature fluctuations that may occur in the wild^29^ (Figure 1B), which has been successfully used to induce breeding in labs studying *A. mexicanus*^30–32^. More specifically, the water temperature was altered via a programmable logic controller (V350-35-T2, Unitronics) to gradually raise up to 26°C and maintained for 48 hours. After this period, the temperature was gradually lowered to 23°C over the course of a few days. Each breeding rack went through a breeding cycle every two or four weeks. During the 48-hour period at 26°C, traps were placed in the tanks to collect embryos twice per week. The embryos were then raised in embryo media containing 0.5X E2 (ZIRC) Media (pH 7.2) with 0.5 mg/L Methylene Blue for less than 10 days in the embryo incubator.

### Gamete collection – *in vitro* fertilization (IVF)

Fish were anesthetized using chilled fish-ready water between 0-4°C. Under anesthesia, female gametes were collected by gently squeezing laterally against the sides of the coelomic cavity while rolling fingers slowly in the direction of the urogenital opening^99^. Ova were collected on a disposable spatula (80881-188, VWR) and placed in a humidified petri dish. Male gametes were expressed from the fish in the same manner as the female gametes. When expressing the male gametes, a capillary tube (2-000-010, Drummond Scientific Company) was placed outside the urogenital opening, and the milt was collected through gentle suction from an aspirator tube (2-000-000, Drummond Scientific Company), then dispensed into a centrifuge tube containing Sperm Extender E400 (ZIRC). For fertilization, the milt solution was dispensed onto the ova, and fish-ready water was added to the clutch to activate the gametes.

### Sperm motility test

Sperm were collected as described above during breeding cycles. The collected milt was diluted at a ratio of 1:10 with Sperm Extender E400 (ZIRC) and stored on ice. The motility of the milt was assessed using Computer-Assisted Sperm Analysis (CASA). Milt-E400 solution was further mixed with fish-ready water at ratios ranging from 1:20 to 1:80 in a PCR tube to activate the milt. 4 µl of the activated milt-E400 solution were then loaded into a chamber of a Leja 20 µm slide (SC 20-01-04 B, Leja). Images of four to six random fields of the slide were captured by Hamilton Thorne Zeiss Axio Lab.A1 microscope and analyzed using the Hamilton Thorne CASA II software (Version: 1.11.9).

### Brightfield imaging of embryos

Embryos were collected in limited embryo media for imaging. Images were captured by LEICA M205 C stereo microscope coupled with Leica Application Suite X software at the 20X to 50X magnification.

### Starvation and breeding experiment

Two independent, age-matched clutches of surface fish and Pachón were selected for this experiment. Only individuals with a good breeding history were chosen. Each clutch was divided into four experimental groups, namely surface fish control, surface fish starvation, Pachón control and Pachón starvation. Breeding attempts were conducted twice a month in the same week. Males paired with females from starvation groups were individually tagged, placed in the starvation tank one day prior to breeding attempts, and returned to regular feeding the day after. Monthly weights were recorded for all fish. For the first clutch, fish were approximately two years old at the beginning of the experiments.

Starvation period started after recording baseline breeding activities recording at 0 month (regular feeding). After two months of starvation, starved fish were returned to regular feeding for a two-month recovery period, followed by another two-month starvation period. We noticed that ovaries exhibited a more liquid-like consistency during breeding cycle, making dissection and collection challenging. Therefore, we waited an additional two weeks before euthanizing the females for tissue dissection. The second clutch consisted of fish around 1 year and 4 months old at the beginning of the experiments. Similar to the first clutch, starvation began after baseline breeding activities at 0 month (regular feeding), followed by two months of starvation. Two weeks later, females were euthanized for tissue dissection. Ovaries were collected, snap frozen in liquid nitrogen and stored at −80°C prior to RNA extraction.

Three breeding cycles were recorded across both clutches. For analysis, all data from control and data from 0 month in starvation groups were used as the control condition, while data from starvation groups at 1 and 2 months after starvation were considered as the starvation condition. Two months of refeeding data from the first clutch were excluded from analysis, though it’s worth noting that, to a lesser extent, starved surface fish resumed some breeding activity during refeeding.

The 2-month starvation experiment of age-matched surface fish and Tinaja (shown in Supplementary Figure 5K) was done in a separate batch of fish. Each experimental group was composed of three female fish. No breeding and spawning were attempted. Similarly, ovaries were collected following dissections, snap frozen in liquid nitrogen and stored at −80°C prior to RNA extraction.

### Sample preparation, LC-MS/MS and data analysis for proteomics

2-cell stage embryos were collected through induced spawning event or IVF. Six biological replicates of each fish morphotype were collected from 6 female fish, and each sample was composed of around 40 embryos. Proteins and lipids were extracted using a Lipid Extraction Kit (ab211044, Abcam) and 1.5 ml BioMasher^®^ II Micro Tissue Homogenizers (749625-0010, DWK Life Sciences) following manufacture protocols. Briefly, embryos were homogenized in 500 μl Extraction Buffer on ice for 1 minute (min), followed by incubation at room temperature for 20 min at 1000 rpm. The homogenate was centrifuged at 10,000 × g for 5 min at 4°C, separating the supernatant and non-polar pellet fractions. The non-polar pellets were saved for proteomic analysis, and the supernatant fraction was used for lipidomic analysis.

The pellet from every two biological replicates were combined for proteomics analysis, resulting in three samples per fish population. Each pellet was resuspended in 100 μl of 8 M Urea with 100 mM of Tris, pH 8.5 for digestion. Samples were reduced with tris(2-carboxyethyl)phosphine (TCEP) to a final concentration of 5 mM and incubated for 30 min at room temperature. To carboxymethylate reduced cysteine residues, 2-chloroacetamide (CAM) were added to a final concentration of 10 mM and incubated for 30 min at room temperature in the dark. Mass spectrometry grade Endoproteinase Lys-C (Promega) was added at a 1:1000 w/w ratio to digest the samples overnight at 37°C. The reactions were subsequently diluted to 2 M Urea by adding 100 mM of Tris, pH 8.5. Then mass spectrometry grade trypsin (V511A, Promega) was added at 1:200 w/w ratio for a second overnight digestion at 37°C. Post-digestion, the samples were centrifuged at 16000 x g for 30 min, and the supernatants were transferred to new tubes. All samples were cleaned by Pierce™ Peptide Desalting Spin Columns (89851, Thermo Fisher Scientific) before quantification using a colorimetric peptide assay (23275, Thermo Fisher Scientific).

Peptide labeling was performed following manufactural protocols with minor modifications. Briefly, 20 µl of anhydrous acetonitrile (ACN) was added to each 0.8 mg TMT label reagent (90061, Thermo Fisher Scientific). The TMT reagents were dissolved after occasional vortexing for 5 min. For each sample, 30 μg of peptides (quantified using the colorimetric peptide assay) were adjusted to 100 µl with 100 mM Tetraethylammonium bromide (TEAB) and mixed with the 20 µl of the respective TMT reagent. Three replicates for surface fish, Pachón and Molino were labeled with TMTpro-126, −127N, −127C; −128N, - 128C, −129N; and −129C, −130C −130C, respectively, and incubated for 1 hour at room temperature. Labeling efficiency was assessed by analyzing 500 ng of each sample via LC-MS/MS using a 2-hour C18 reverse-phase (RP) gradient on an Orbitrap Eclipse Tribrid Mass Spectrometer (Thermo Fisher Scientific) with a FAIMS Pro interface, equipped with a Nanospray Flex Ion Source, and coupled to a Vanquish Neo System. The TMT labeling efficiency exceeded 99% for all detected peptides (data not shown). The 9 differentially labeled samples (40 µl each) were combined in a single tube and the resulting volume was reduced to less than 10 µl using a SpeedVac concentrator (Savant). High pH RP fractionation was then performed following manufactural protocols (84868, Thermo Fisher Scientific) to prepare the samples for LC-MS/MS analysis.

TMT-labeled peptides were analyzed as described above for labeling efficiency testing. Peptides (22 µl for each HpH RP fraction) were loaded on an Acclaim PepMap 100 C18 trap cartridge (0.3 mm inner diameter (i.d.), 5 mm length (L); Thermo Fisher Scientific) with the Neo loading pump at 2 µL/minute via the autosampler. A 75 µm i.d. analytical microcapillary column was packed in-house with 250 mm of 1.9 μm ReproSil-Pur C18-AQ resin (Dr. Masch). AgileSLEEVE (Analytical Sales & Products) was used to maintain column temperature at 40°C. The organic solvent solutions consisted of water:ACN:formic acid at 95:5:0.1 (v/v/v) for buffer A (pH 2.6) and 20:80:0.1 (v/v/v) for buffer B. Chromatography was performed using the following gradient: a 5-min column equilibration at 1% buffer B; a 74-min ramp to reach 30% buffer B; a 20-min ramp from 30% to 60% buffer B; a 3-min increase to reach 90% buffer B; a 10-min wash at 90% buffer B; a 0.1-min return to 1% buffer B; and a final 12-min column re-equilibration at 1% buffer B. The Neo pump flow rate was set to 0.180 μl/min.

The Orbitrap Eclipse was set up to run the TMTpro-SPS-MS3 method. Briefly, peptides were scanned from 400-1600 m/z in the Orbitrap at 120,000 resolving power before MS2 fragmentation by collision-induced dissociation (CID) at 35% normalized collision energy (NCE), and detection in the ion trap set to turbo detection. Dynamic exclusion was enabled for 45 seconds. Carbamidomethyl (+57.0215 Da on cysteine) and TMT10plex (+229.163 Da on lysine and peptide N-termini) were searched statically, while methionine oxidation (+15.9949 Da) was searched as a variable modification. Synchronous precursor scanning (SPS) was used to select the top 10 MS2 peptides for TMT reporter ion detection in the Orbitrap using HCD fragmentation at 65% NCE at resolving power of 50,000.

The LC/MS dataset was processed using Proteome Discoverer 3.0 (Thermo Fisher Scientific). MS/MS spectra were searched against the *A. mexicanus* protein database (NCBI 2022-07) complemented with common contaminants. SEQUEST-HT implemented through Proteome Discoverer was set up as: precursor ion mass tolerance 10 ppm, fragment mass tolerance 0.6 Dalton, up to two missed cleavage sites, static modification of cysteine (+57.021 Da), and lysine and peptide N-termini with TMT tag (+229.163 Da) and dynamic oxidation of methionine (+15.995 Da). Results were filtered to a 1% FDR at peptides levels using Percolator through Proteome Discoverer. MS3 spectra were processed to extract intensity for each reporter ion.

Protein mass spectrometry data was filtered to include only proteins detected in all 9 samples. Samples were normalized using NormalyzerDE v1.20.0^44^, and median normalization was chosen for further downstream analysis. As Pachón samples were collected using different methods, a Mann-Whitney U test was performed within these samples, confirming that collection method had no significant effect on the proteomics result. Protein accession IDs were mapped to NCBI gene IDs through a combination of rentrez v1.2.3^100^, biomaRt v2.58.2^101^ and manual NCBI searches. Fold changes for each protein were calculated using limma v3.58.1^45^ and resulting p values were adjusted with Benjamini-Hochberg method. The significance threshold was set with an absolute average log2 fold change greater than 0.5 and an adjusted p-value smaller than 0.05. Analysis was done in R v4.3.3 available at https://github.com/Xiazistarry/cavefish.

### Sample preparation, LC-MS/MS and data analysis for lipidomics

Lipid extraction was described as above. The supernatant fraction of 6 individual biological replicates were transferred into new tubes, dried and then resuspended in 16 μl of lipid sample buffer (Butanol: Isopropanol:water in a 8:23:69 ratio (v/v/v), and 5 mM phosphoric acid). Additionally, three technical replicates of embryo water or sample buffer-only samples were included as negative controls. Using gel loading tips, 15 µl of each sample was transferred into glass sample vials, 5 µl each was analyzed in negative and positive ionization modes per sample.

A Q Exactive^TM^ Plus Hybrid Quadrupole-Orbitrap^TM^ Mass Spectrometer (Thermo Fisher Scientific) coupled with an UltiMate 3000 RSLCnano HPLC system was used for LC-MS/MS analysis. A 35-min chromatographic separation was carried out using a Acquity UPLC^®^ CSH^TM^ C18 (186005297, Waters Corporation) column with 1.7 μm particles and 2.1 mm × 100 mm (i.d./L). Mobile Phase A was ACN:H_2_O at a ratio of 60:40% (v/v), 0.1% formic acid, and 10 mM ammonium formate. Mobile Phase B was Isopropanol:ACN at a ratio of 90:10% (v/v), 0.1% formic acid, and 10 mM ammonium formate. The flow gradient condition was 30% solvent B to 100% solvent B over 18 min, 6 min at 100% solvent B, a return to 30% solvent B over 0.1 min then followed by a 12-min re-equilibration at 30% solvent B prior to the next injection. The flow rate was maintained at 100 μl/min.

A Q Exactive Plus™ Mass Spectrometer (QE+) operating in positive or negative ion modes with a scan range of 200-1500 m/z, 3.3 kV (−) or 3.8 kV (+) spray voltage and 300°C ion transfer tube temperature. The precursor ion was detected in the orbitrap at 140,000 resolution with an AGC target of 3e6 and 100 ms maximum injection time. The product ion was isolated in a quadrupole with NCE at 17-30-50% higher-energy collisional dissociation (HCD) collision energy and detected in the orbitrap at 17,500 resolution. The AGC target for the MS/MS ions was set at 1e5 with a maximum injection time of 100 ms. Each sample was analyzed once in positive and negative ion modes independently. MS and MS/MS datasets were interpreted using Thermo Xcalibur Qual Browser and Compound Discoverer v3.3.

Lipid mass spectrometry data was filtered to include only lipids that were detected in all samples, had valid name and formula, passed false background filtering, featured annotated delta masses between −10 and 10, and had average abundances in negative controls that were less than five times the average abundance in actual samples. Counts for the same lipid names were consolidated. Outlier samples, including two surface fish, one Pachón and one Molino, were excluded based on significantly deviating total abundances. Similar to proteomics data analysis, the lipidomics data was normalized using NormalyzerDE v1.20.0^44^, and median normalization was selected for further downstream analysis. As Pachón samples were collected using different methods, a Mann-Whitney U test was performed within these samples, confirming that collection method had no significant effect on the lipidomics results in positive ion mode. Lipids were categorized using a combination of RefMet v1.0.0^49^ and manual annotation on LIPID MAPS^® 50–52^ (https://www.lipidmaps.org) and PubChem^53^ (https://pubchem.ncbi.nlm.nih.gov). Fold changes for each protein were calculated using limma v3.58.1^45^ and resulting p values were adjusted with Benjamini-Hochberg method. The significance threshold was set with an absolute average log2 fold change greater than 0.5 and an adjusted p-value smaller than 0.05. Analysis was done in R v4.3.3 available at https://github.com/Xiazistarry/cavefish.

### Whole ovary dissociation for scRNA-Seq and stage-specific bulk RNA-seq

Dissociation methods for different stages of surface fish and cavefish ovary cells were modified from zebrafish ovary cell dissociation protocols^71,83,102^. The digestive enzyme mixture is composed of 3 mg/ml collagenase I (C0130, Sigma-Aldrich), 3 mg/ml collagenase II (C6885, Sigma-Aldrich) and 1.6 mg/ml hyaluronidase (H4272, Sigma-Aldrich) in L15 media (112-029-101, Quality Biological). Fish were fasted one day before cell dissociation. Each sample and library were collected and constructed from a single fish, and the fish being used were aged-matched.

The left side ovary from each adult fish was dissected and stored on ice in an RNA Lobind tube (0030108310, Eppendorf north america) containing 3 ml of L15 for no more than 40 min. Tissues were minced with dissection scissors into pieces. L15 media were replaced with 3 ml of digestive enzyme mixture after the tissue settled down to the bottom of the tube. Samples were incubated on an orbitron rotator (260200, Boekel Scientific) at room temperature until no obvious cell clumps were observed (35∼45 min, depending on samples). Cells were centrifuged at 300 g for 3 min at 4°C, resuspended in 3 ml of TrypLE (12604021, Thermo Fisher Scientific) and incubated on the orbitron rotator at room temperature for 15 min. 3 ml of 20% FBS (SH30071.03E, Cytiva) in L15 were added to the cell suspensions and incubated on the orbitron rotator at room temperature for 1 min to stop TrypLE reaction. Cell suspensions were then added with 15 ml of L15 and centrifuged at 300 g for 3 min at 4°C to get the cell pellets. The cell pellets were resuspended and washed three times with 5 ml of L15 using a P1000 pipette and centrifuged at 300 g for 3 min. Cell pellets were resuspended in 2 ml L15 and filtered through a 70 µm nylon filter (352350, Corning) and then through a 40 µm nylon filter (352340, Corning) into Lobind tubes to remove large cells and cell clumps. The filters were washed twice with 2 ml L15. The filtrates were centrifuged at 300 g for 3 min and resuspended in 1 ml of 2% BSA (A9418, Sigma-Aldrich) in phosphate-buffered saline (PBS) (20012050, Thermo Fisher Scientific) and transferred into Lobind tubes.

The right side ovary from each adult fish were dissected and stored on ice in an RNA Lobind tube containing 2 ml of L15 for no more than 40 min. L15 media were replaced with 2 ml of digestive enzyme mixture. Samples were incubated on the orbitron rotator at room temperature for 12∼15 min. Cell suspensions were then filtered through a series of 300 um cell strainer (43-50300-03, pluriSelect), 100 um cell strainer (10199-658, VWR international) and 40 µm cell strainer to RNA Lobind tubes to separate ovary cells at sizes > 300 µm, 100∼300 µm, 40∼100 µm and < 40 µm. Tubes and filters were washed twice with 2 ml L15. 20 ml of fresh L15 were added to the cell suspensions containing oocytes and somatic cells under 40 µm to dilute the digestive enzyme mixture. 5 ml of fresh L15 were used twice or more to wash and collect the oocytes on the filter. Different sizes of ovary cells were then incubated in L15 for an hour at room temperature to let oocytes settle down at the bottom of the tube. For oocytes < 100 µm, cell suspensions were centrifuged at 800 rpm for 2 min to collect cell pellets. For oocytes > 100 µm, supernatant L15 were removed without centrifugation. Cell pellets were snap frozen in liquid nitrogen and stored at −80°C prior to RNA extraction.

### Single cell library construction and sequencing

scRNA-seq libraries were prepared by Stowers Sequencing core. Dissociated, sorted cells were assessed for concentration and viability using a Luna-FL cell counter (Logos Biosystems). Cells deemed to be at least 84% viable were loaded based on live cell concentration onto a Chromium iX Series instrument (10x Genomics) and libraries prepared using the Chromium Next GEM Single Cell 3’ Reagent Kits v3.1 (10x Genomics) according to manufacturer’s directions. Resulting cDNA and short fragment libraries were checked for quality and quantity using a 2100 Bioanalyzer (Agilent Technologies) and Qubit 2.0 Fluorometer (Thermo Fisher Scientific). With cells captured estimated at ∼1,100-5,500 cells per sample, libraries were pooled and sequenced to a depth necessary to achieve at least 38,000 mean reads per cell on an Illumina NextSeq 2000 instrument utilizing RTA and instrument software versions current at the time of processing with the following paired read lengths 28*10*10*90bp.

### scRNA-Seq analysis

10x scRNA-Seq data was analyzed using Cell Ranger v7.1.0. Raw reads were demultiplexed into FASTQ files with cellranger mkfastq. The Cell Ranger count pipeline was used to align reads, filter low-quality sequences, and perform barcode and UMI counting. Alignments were conducted against the Astyanax_mexicanus-2.0 genome, using Ensembl Ens_110 annotation. CellBender v0.3.0^103^ was applied to mitigate ambient RNA effects from over digested later-stage oocytes. For quality control, cells with a total RNA count (nCount_RNA) above 500 were retained for downstream analysis using Seurat v5.1.0^104^ in R 4.3.1. Cells with unique RNA feature counts under 200 or above three standard deviations from the mean were classified as low quality and excluded. DoubletFinder v2.0.4^105^ was used to identify and remove doublets. After quality control, 1344∼5037 single cells were detected per sample. Each of the two samples in each fish population was normalized individually using the SCTransform workflow, followed by merging into a combined sample per population. The merged samples were integrated across populations with Harmony v1.2.0^106^ to address batch effects. We selected the top 3,000 variable genes for PCA, retaining the first 40 principal components (PCs) based on the elbow plot for downstream UMAP and cell clustering analyses. Cell clusters were annotated according to the expression of established cell markers. Cell proportion analysis was performed using scCustomize v1.1.3^76^ and scProportionTest v0.0.0.9000^77^. For targeted analysis, follicular somatic cells were subsetted and processed using the same approach. Monocle3 v1.3.4^107^ was used to perform trajectory inference, with the expression matrix extracted from the Seurat object as input. Cells expressing *lhx9* were designated as the root cells of the trajectory, from which pseudo-time was inferred. DEGs between surface fish and cavefish across all cell types were identified using the FindMarkers function in Seurat v5.1.0 using the default Wilcoxon Rank Sum test (Mann-Whitney U test). The significance threshold was set with an absolute average log2 fold change greater than 0.5 and an adjusted p-value smaller than 0.05. GO enrichment analysis on DEGs in follicular somatic cells was conducted through web-based tool ShinyGO v0.81^80,81^, using Ensembl IDs of DEGs as search input and common genes existed in all populations after quality control as the background. Analysis was done in R v4.3.1 available at https://github.com/Evenlyeven/cavefish_ovary_scRNAseq.

### Total RNA extraction, library construction and sequencing for Bulk RNA-seq

Total RNA was extracted using standard phenol/chloroform extraction. 1 ml of Trizol (15596026, Thermo Fisher Scientific) was used per 50 to 100 mg tissue. Samples were homogenized with Trizol and 3.0 mm TriplePure M-Bio Grade beads (D1132-30, Benchmark Scientific) on Beadbug 6 homogenizer (D1036, Benchmark Scientific). After a brief centrifugation, 200 µl of chloroform was added per 1 ml of Trizol, then vortexed for 15 seconds and incubated at room temperature for 2 min. Samples were centrifuged at 12,000 g for 15 min at 4°C. Upper transparent aqueous phase was transferred carefully without disturbing the interphase into a fresh tube and 500 µl of Isopropyl alcohol per 1 ml of Trizol was added. 2 µl of GlycoBlue Coprecipitant (AM9515, Thermo Fisher Scientific) was added to visualize RNA pellets. Samples were incubated at room temperature for 10 min and centrifuged at 12,000 g for 10 min at 4°C. Supernatant was removed and RNA pellets were washed twice with 1 ml of 75% ethanol per 1 ml Trizol used initially. Samples were mixed by vortexing and centrifuged at 7400 g for 5 min at 4°C. RNA pellets were air dried for 15 min and dissolved in RNase free water. RNA solutions then went through on column purification and DNA digestion using RNeasy Mini Kit (74104, QIAGEN INC) and DNase kit (79254, QIAGEN INC).

Total RNA-Seq libraries were generated from 3.8-500 ng of high-quality total RNA, as assessed using the Bioanalyzer (Agilent), according to the manufacturer’s directions using a 200-5-fold dilution of the universal adaptor and 16-9 cycles of PCR per the respective input mass with the NEBNext Ultra II Directional RNA Library Prep Kit for Illumina (E7760L, NEB), the NEBNext rRNA Depletion Kit v2 (Human/Mouse/Rat) (E7400L, NEB), and the NEBNext Multiplex Oligos for Illumina (96 Unique Dual Index Primer Pairs) (E6440S, NEB) and purified using the SPRIselect bead-based reagent (B23318, Beckman Coulter). Resulting short fragment libraries were checked for quality and quantity using the Bioanalyzer and Qubit Fluorometer (Life Technologies). Equal molar libraries were pooled, quantified, and converted to process on the Singular Genomics G4 with the SG Library Compatibility Kit, following the “Adapting Libraries for the G4 – Retaining Original Indices” protocol. The converted pools were sequenced to the required number of reads per library on F2 flow cells (700102) or F3 flow cells (700126) on the G4 instrument with the PP1 and PP2 custom index primers included in the SG Library Compatibility Kit (700141), using Instrument Control Software 23.08.1-1 with the following read lengths: 8 bp Index1, 100 bp Read1, 8 bp Index2, and 100 bp Read2. Following sequencing, sgdemux 1.2.0 was run to demultiplex reads for all libraries and generate FASTQ files.

### Bulk RNA-seq analysis

Raw reads were demultiplexed into Fastq format allowing up to one mismatch using Illumina bcl-convert 3.10.5. Reads were aligned to UCSC genome Astyanax_mexicanus-2.0 with STAR aligner v2.7.10b^108^, using Ens_110 gene models. TPM values were generated using RSEM v1.3.1^109^. The STAR count tables were used for differential gene expression analysis using edgeR v3.42.4^110^ in R 4.3.1. Lowly expressed genes were filtered out with filterByExpr’s default settings. All samples from each experiment were sequenced within the same batch and normalized together using Trimmed Mean of M-values (TMM) normalization. PCA was conducted using prcomp in ggfortify v0.4.17^111^ and plotted using autoplot in ggplot2 v3.5.1^112^. Differential gene expression analysis and comparisons were performed between the same cell categories between surface fish and cavefish and different conditions in the same population. The significance threshold was set with an absolute average log2 fold change greater than 0.5 and an adjusted p-value smaller than 0.05. GO enrichment analysis on DEGs was performed through web-based tool ShinyGO v0.81^80,81^, using Ensembl IDs of DEGs as search input and all genes after low expression filtering as the background. Analysis was done in R v4.3.1 available at https://github.com/Xiazistarry/cavefish.

### Ovary fixation, sectioning and staining for light and electron microscopy imaging

For H&E staining, the procedure was adapted from zebrafish ovary protocols^113^ and lab notes from previous lab members^22^. Specifically, dissected ovaries were fixed in Davidson’s fixative overnight at room temperature without shaking. The fixed samples were washed 3 times with 70% ethanol and then paraffin processed and embedded. Tissues were sectioned at 5 μm and slides were dried in a 60°C oven for 20 min. Slides were deparaffinized, then stained with hematoxylin for 2 min and eosin for 30 seconds. Slides were dehydrated and placed in xylene for coverslipping with Cytoseal. Images were obtained using a VS120 virtual slide microscope (Olympus).

For electron microscopy imaging, ovaries were dissected from around 500 dpf age-matched surface fish, Pachón and Molino. Sections of ovaries less than 1 mm^3^ were fixed in 2.5% gluteraldehyde, with 2% paraformaldehyde, 1 mM CaCl_2_, 50 mM sodium cacodylate and 1% sucrose at room temperature with light agitation for 1 hour followed by overnight at 4°C. After 4 x 15 min buffer rinses, a secondary fix of 50 mM buffered 2% osmium tetroxide and 1% potassium ferricyanide was performed for 2 hours. 4 × 10 min buffer rinses followed by 4 × 10 min water rinses preceded a 0.5% aqueous uranyl acetate incubation overnight at 4°C. The next morning samples were washed 4 x 15 min in water then sequentially dehydrated in increasing concentrations of ethanol over 2 hours with 3 × 100% ethanol washes at the end. Samples were infiltrated with Hard Plus Epon without accelerator at 25, 50, 75, and two changes of 100% resin for at least two hours each step with the final exchange over 48 hours at 4°C. Finally, 4 exchanges of 100% resin with accelerator were performed using a BiowavePro+ microwave (Pelco) at 250W for 3 min each. Samples were embedded and polymerized for 48 hours at 60°C and were cut at 80 nm onto slides using a Leica Artos 3D Ultramicrotome and then stained with Sato’s Triple lead, 1% aqueous uranyl acetate and Sato’s Triple Lead again for 10 minutes at room temperature for each step. The resulting slides were coated with 4 nm carbon using a Leica Ace600 sputter coater. Samples were imaged using a Zeiss Merlin SEM with a BSD4 detector at 8.0 kV and 700 pA.

### Oocyte staging and size measurement for H&E-stained ovary images

H&E-stained ovary images vary wildly in sizes. Therefore, they were all padded to a consistent 10,000 X 10,000 canvas size. Oocyte staging was performed based on established criteria from zebrafish^64,114^. Six images, with hundreds of examples of stage I, stage II, and stage III and later were used to train a YOLOv8 object detector. This classifier was then applied to each padded image from the datasets. Areas and counts for the three stages were compiled and the results plotted. All analysis was done in python using jupyter notebooks available at https://github.com/jouyun/Rohner_2025_Ovary.

### Statistical analysis

All statistical analysis were conducted in R v4.3.1 or v4.3.3. Data distribution was checked before choosing specific statistical test. All statistical tests were mentioned in the figure legends accordingly,

## Data availability

Protein mass spectrometry data are available at the ProteomeXchange Consortium with the dataset identifier PXD058862 via the MassIVE repository. Lipid mass spectrometry data are available at MSV000096792 via the MassIVE repository. Single cell and bulk RNA sequencing data are deposited into GEO SuperSeries GSE284736. Interactive data for single cell RNA sequencing is available on ShinyApp https://simrcompbio.shinyapps.io/AstMex_ovary_scRNAseq/.

## Acknowledgement

This work was supported by institutional funding, NIH New Innovator Award 1DP2AG071466-01, and NIH grant R24OD030214 to NR. Cavefish cartoon illustrations were made by Mark Miller, and others were made on BioRender. We would also like to thank members of the Stowers cavefish team – Adam Potter, David Jewell, Molly Miller, Cole Biesemeyer and Seth Bodtke for their support in fish husbandry and care. Our thanks also go to current and past members of the Rohner lab, co-advisor Dr. Paul Trainor, and to our graduate school committee members, Dr. Matthew Gibson, Dr. Kausik Si, and Dr. Jennifer Gerton, for their valuable feedback on this project. We gratefully acknowledge the technical support provided by the Stowers core facilities and students, particularly Selene Swanson, Yan Hao and Dr. Laurence Florens for proteomics and lipidomics sample preparation and data acquisition; Dr. Joseph Varberg, Dr. Hua Li and Gabriel da Silva Pescador for the general inputs for proteomics and lipidomics data analysis; Hannah Wilson for preparing ovary histological sections and performing H&E staining; Cindy Maddera for scanning H&E-stained images; Xia Zhao for preparing ovary fixation and sections for electron microscopy staining; Kaitlyn Petentler, Allison Scott and Kate Hall for scRNA-seq library construction; Sean McGrath, Michael Peterson, Anoja Perera and Amanda Lawlor for bulk RNA-seq sample optimization, library construction and sequencing; and Dr. Lu Deng for suggestions on scRNA-seq analysis. Special thanks to Dr. Yann Gibert from University of Mississippi Medical Center for sharing protocols for lipidomics in fish embryos; Dr. Bruce W. Draper from University of California, Davis, for sharing protocols on single-cell dissociation in fish ovaries and general input on scRNA-seq analysis; and Dr. Kellee R. Siegfried from University of Massachusetts Boston for her assistance with ovary histological sample preparation.

## Author contribution

F.X. and N.R. conceived the project as part of F.X.’s thesis research to fulfill the requirements for the Graduate School of the Stowers Institute for Medical Research. F.X. and N.R. designed the experiments, which were performed by F.X., A.S., E.F., P.M., and S.N. Data analysis was performed by F.X., D.W., S.B. and S.M. The manuscript was prepared by F.X. and N.R. with input from all authors.

## Declaration of interests

All authors declare no competing interests.

---

**Supplementary Figure 1.**
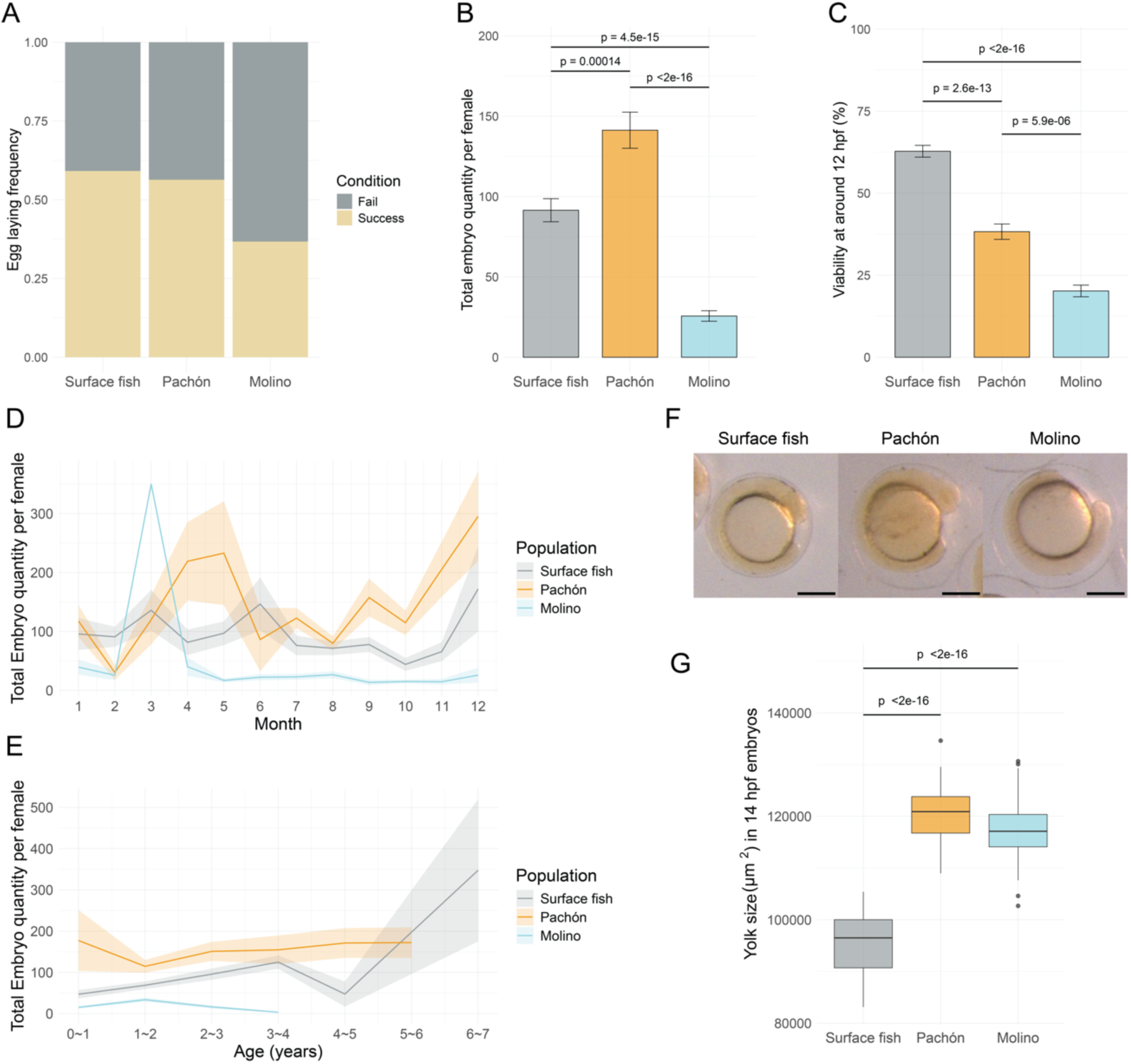
Independent cavefish populations have unique reproductive output compared to surface fish. (A-C) The measurement of egg laying frequency, clutch size and embryo viability at the time of collection from one year of estimated reproductive recording. Egg laying frequency (A) was calculated from all spawning attempts (n=394, surface fish; n=291, Pachón; n=381, Molino). Clutch size (B) and embryo viability (C) were measured from successful spawning events (spawned eggs > 0; n=233, surface fish; n=164, Pachón; n=140, Molino). Kruskal-Wallis Test was performed for statistically analysis for the comparisons of clutch size (Kruskal-Wallis chi-squared = 125.14, df = 2, p-value < 2.2e-16) and embryo viability (Kruskal-Wallis chi-squared = 157.83, df = 2, p-value < 2.2e-16) among three populations. Pairwise comparisons were performed by Dunn’s test. P-values were adjusted with Bonferroni method. (D) Egg production seasonality in surface fish, Pachón and Molino in the lab. It was measured from successful spawning events. (E) Egg production in different ages of surface fish, Pachón and Molino in the lab. It was measured from successful spawning events. (F) Representative images of 14 hpf surface fish, Pachón and Molino embryos. Scale bar, 200 μm. (G) Yolk size measurements in 14 hpf surface fish, Pachón and Molino embryos (n=29, surface fish; n=32, Pachón and n=96, Molino). One way ANOVA was performed for statistically analysis for the comparisons among three populations (F = 197, Pr(>F) < 2e-16). Pairwise comparisons were performed by Tukey’s HSD test.

**Supplementary Figure 2.**
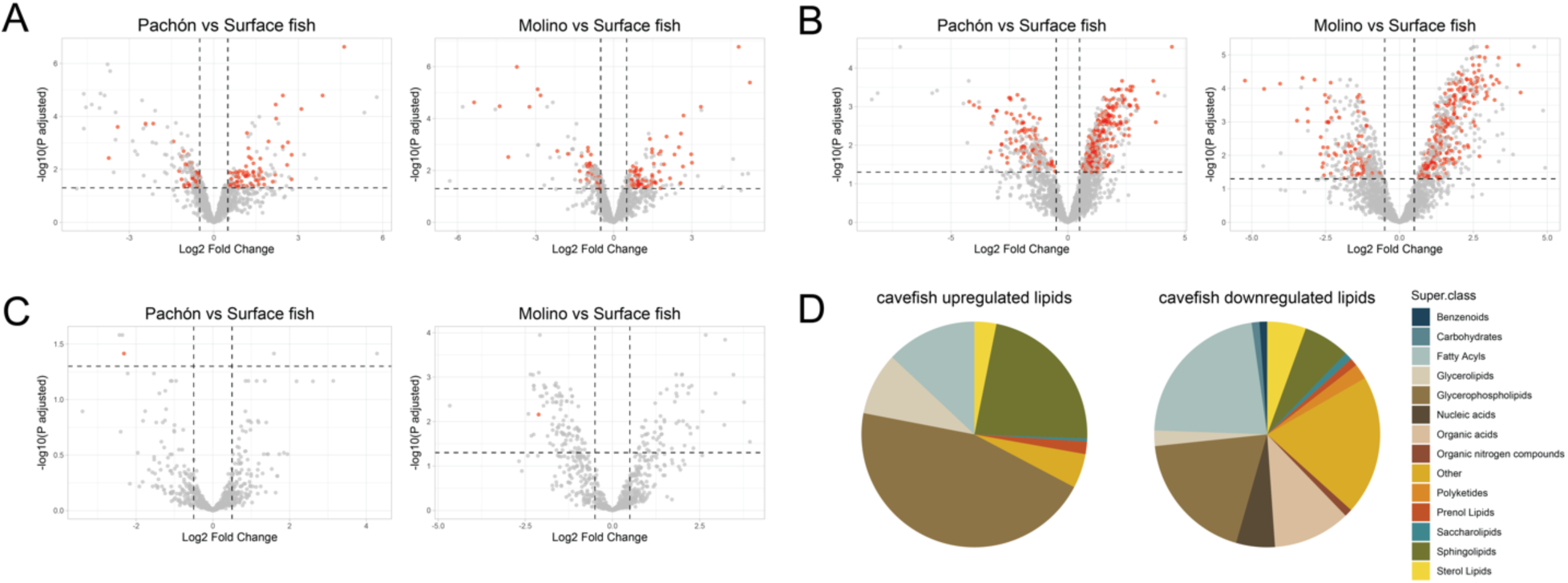
Unique yolk protein and lipid profiles between surface fish and cavefish. (A) Comparisons of protein profiles between surface fish and cavefish 2-cell stages embryos. Commonly enriched proteins in surface fish or cavefish are highlighted in red. (B-C) Comparisons of lipid profiles between surface fish and cavefish 2-cell stages embryos in positive ion mode (B) and negative ion mode (C). Commonly enriched lipid species in surface fish or cavefish are highlighted in red. Vertical dash lines indicate the absolute number of Fold Change equals 0.5; horizontal dash lines indicate the p adjusted value equals 0.05. (D) Lipid categories for upregulated (left) and downregulated (right) lipids in cavefish compared to surface fish from positive ion mode.

**Supplementary Figure 3.**
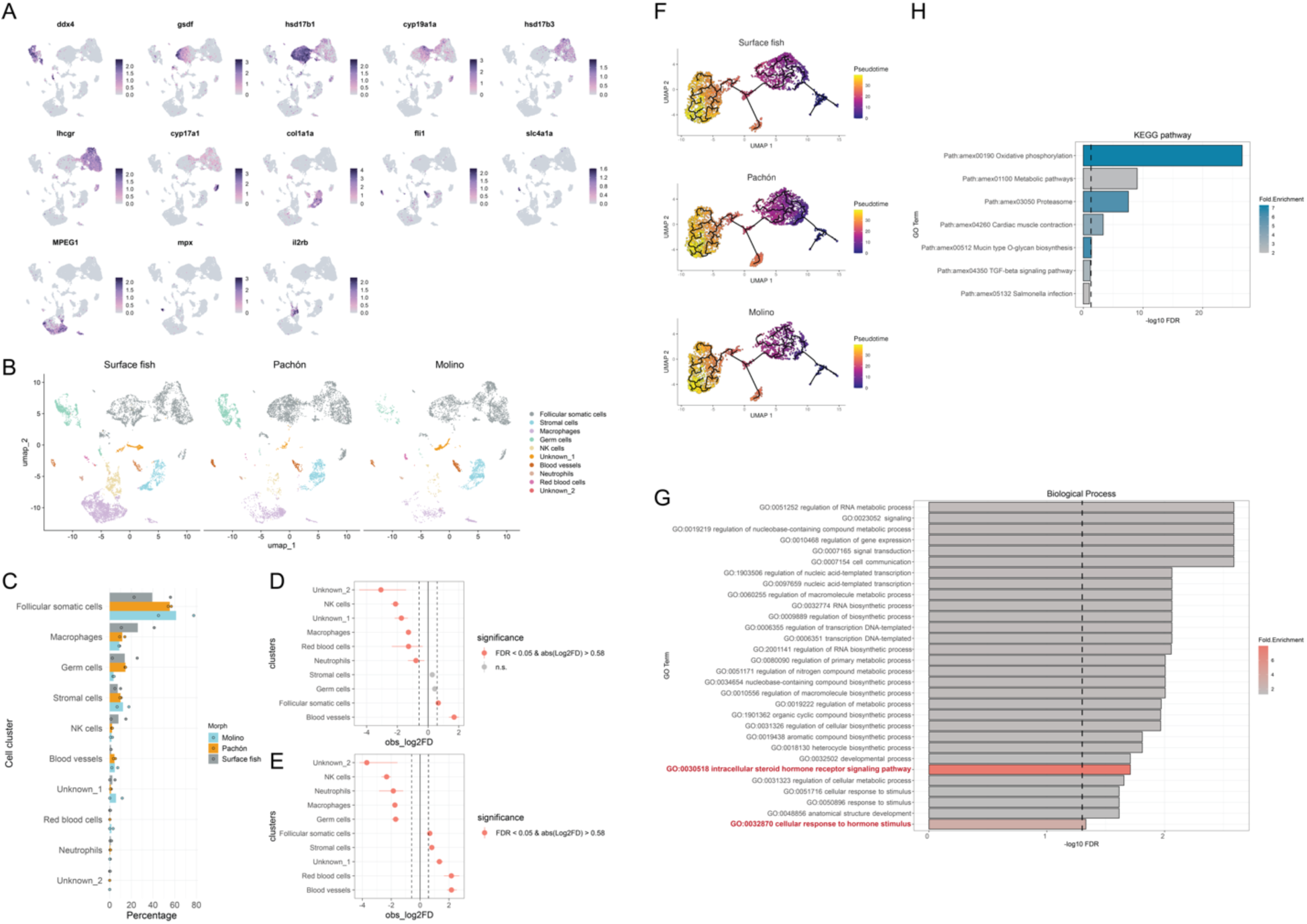
scRNA-seq analysis in *A. mexicanus* reveals an enriched population of follicular somatic cells in cavefish compared to surface fish. (A) Gene expression plots of known cell marker genes label the major cell types shown in Figure 3C. (B) UMAP plots of the adult fish ovaries from surface fish, Pachón and Molino. Cells are color-coded by computationally determined cell clusters. (C) Annotated cell proportions in three fish populations. Each dot represents one single-cell library generated from individual fish. (D-E) Cell proportion analysis by scProportionTest^77^ revealed significantly increased and decreased cell proportions between surface fish and Pachón (D) as well as surface fish and Molino (E). Dash lines indicate the absolute number of Fold Change equals 0.5. (F) Trajectory analysis of follicular somatic cells in three fish populations. The color gradient from dark blue to bright yellow indicates the relative pseudotime during follicular somatic cells development. The starting population was pre-follicle cells labeled by *lhx9* as suggested in zebrafish ovary scRNA-seq analysis^71^. (G) Biological Process GO term analysis of upregulated genes in cavefish follicular somatic cells compared to surface fish. (H) Kyoto Encyclopedia of Genes and Genomes (KEGG) pathway analysis of downregulated genes in cavefish follicular somatic cells compared to surface fish. The dash line indicates significant threshold (FDR = 0.05).

**Supplementary Figure 4.**
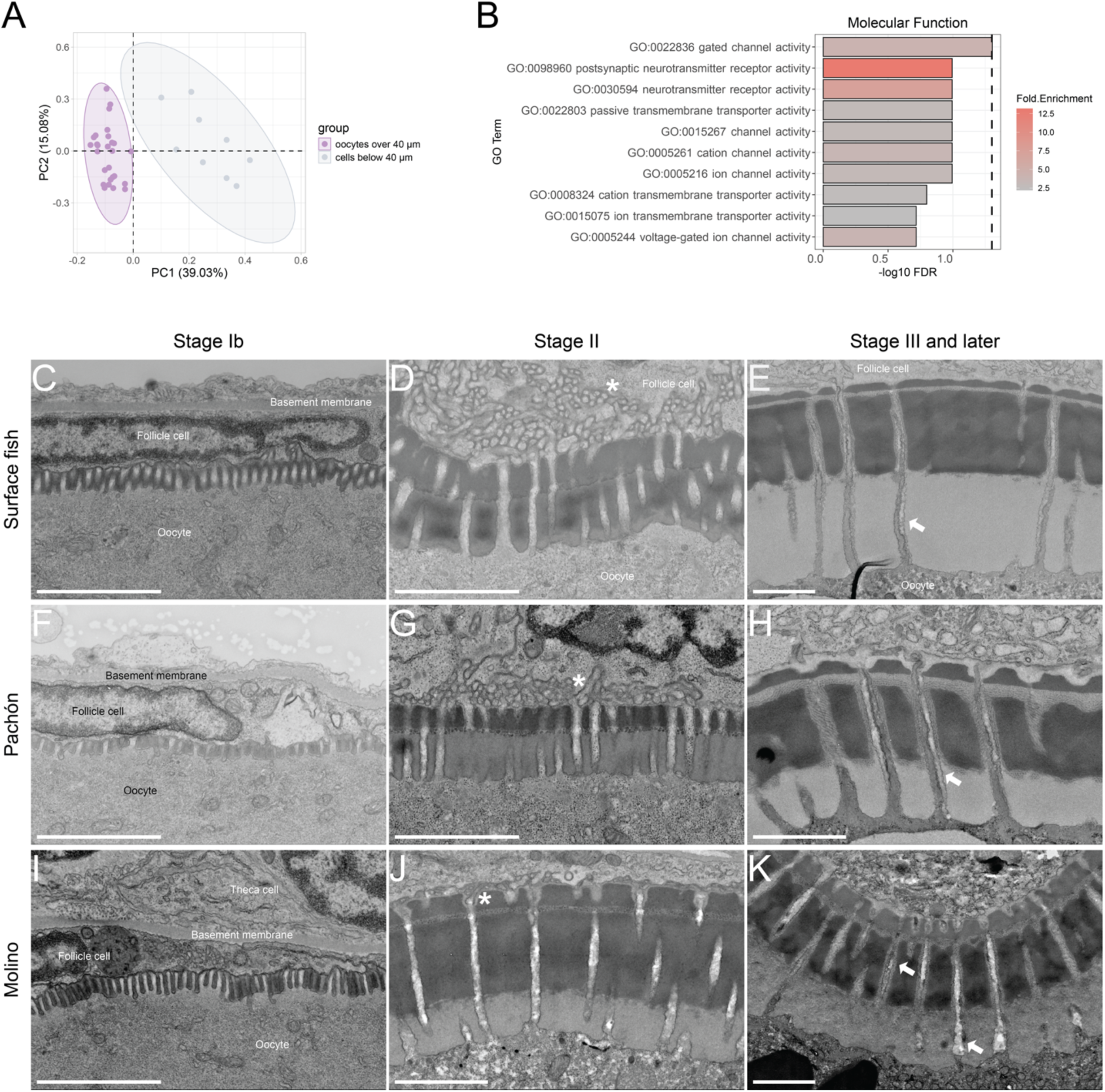
Ovary transcriptomic and morphological characterization in *A. mexicanus*. (A) PCA plot of transcriptomic profiles of cells at different size categories in three fish populations. Cells larger than 40 µm were oocytes, shown in purple. Cells smaller than 40 µm include somatic cells, germ cells and early-stage oocytes, as indicated by grey color. These two groups were transcriptionally distinct to each other. (B) Molecular function GO term analysis of commonly upregulated genes in oocyte size between 100 and 300 µm of both cavefish compared to surface fish. The dash line indicates significant threshold (FDR = 0.05). (C-K) Representative images of VE between follicle cells and oocytes by electron microscopy are shown for (C-E) surface fish, (F-H) Pachón and (I-K) Molino. Scale bar, 2 μm. In all representative images, follicle cells are located above the VE while oocytes are below. In stage Ib oocytes, the VE can vary from 0 to 3 layers depending on developmental stages. Three VE layers were observed in stage II, III and later-stage oocytes. In all three populations, microvilli extending from oocytes were predominantly observed at early stages (C, F and I), with extensions passing through pore canal towards follicle cells (D and G, asterisk). At the end of stage II (J), microvilli from follicle cells extended into the pore canals (J, asterisk), accompanied by the retraction of oocyte microvilli on the follicle cell side, forming two microvilli from both follicle cells and oocytes in the same pore canal to enhance the intercellular interaction as shown in stage III and later stages oocytes (E, H and K, arrows). Follicle cell microvilli can extend to the oocyte surface.

**Supplementary Figure 5.**
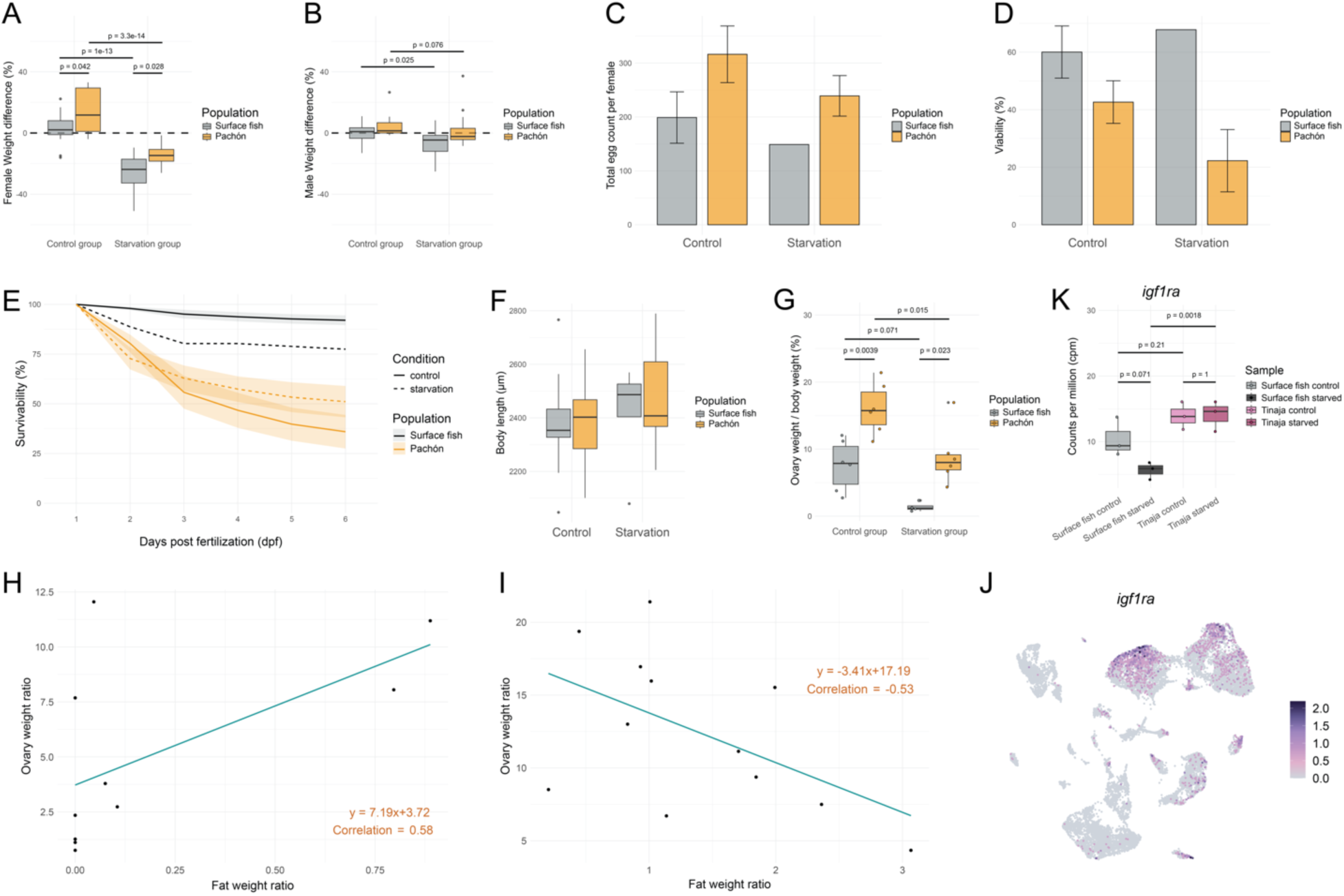
Pachón cavefish reproduction is starvation resistant. (A) Female and (B) male fish weight changes during starvation. Dash line represents no weight difference. Values above the dash line suggest weight gain, and values below dash line represent weight loss. Weight differences were calculated compared to initial weight at 0 month. (A) Both surface fish and Pachón females lost weight during starvation. Weight differences at 1 and 2 months after starvation were combined for simplicity (n=16, surface fish starvation group; n=20 for all others). ART ANOVA was performed for statistically analysis for the comparisons (Group:Population F = 3.2656e-04, Pr(>F) = 0.98563228). Post-hoc tests were performed by built in art.con() function. (B) Surface fish, not Pachón males lost weight significantly during a 5-day starvation. Males paired with starved females were only starved during the breeding weeks when placed in the female tank. Weights were measured at the end of each breeding week. Weight differences at 1 and 2 months were combined for simplicity (n=20 for all four groups). ART ANOVA was performed for statistically analysis for the comparisons (Condition:population F = 0.72357, Pr(>F) = 0.3976490). Post-hoc tests were performed by built in art.con() function. Clutch size (C) and embryo viability (D) were measured from successful spawning events (spawned eggs > 0; n=18, surface fish control; n=16, Pachón control; n=1, surface fish starvation; n=7, Pachón starvation). (E) Offspring survivability was calculated and normalized to the total viable eggs at 1 dpf. (F) Body length of offspring larvae was measured at 6 dpf (up to 5 larvae were selected from the same spawning event; n=61, surface fish control; n=58, Pachón control; n=5, surface fish starvation; n=25, Pachón starvation). ART ANOVA was performed for statistically analysis for the comparisons (Group:Population F = 1.1407e-02, Pr(>F) = 0.915091). Post-hoc tests were performed by built in art.con() function. No pairwise comparison shows statistically significance (p < 0.05). Clutch size, viability, offspring survivability and 6 dpf offspring body length were affected by starvation in both surface fish and Pachón. (G) The ratio of ovary weight to body weight was reduced in both surface fish and Pachón during starvation (n=4, surface fish starvation; n=6 for all other groups). Two way ANOVA was performed for statistically analysis for the comparisons (Group:Population F = 0.092, Pr(>F) = 0.76483). Pairwise comparisons were performed by Tukey’s HSD test. (H) A positive correlation of ovary weight and fat weight is suggested in surface fish. (I) A negative correlation of ovary weight and fat weight is suggested in Pachón. (J) The gene expression plot of *igf1ra* shows its main expression in follicular somatic cells and stromal cells (see cell cluster annotation in Figure 3C). (K) *igf1ra* expression level (cpm) in the ovaries of surface fish and Tinaja control and starvation groups. Adjusted p value was calculated from bulk RNA-seq analysis.

**Supplementary Table 1 List of protein list with commonly increased or reduced abundances in 2-cell stage embryos of both cavefish compared to surface fish.** Blue text represents potential gene names or Ensembl IDs derived from protein IDs. All abundance values were median-normalized and log2 transformed.

## Notes

### Competing Interest Statement

The authors have declared no competing interest.

